# GSearch: Ultra-Fast and Scalable Microbial Genome Search by Combining K-mer Hashing with Hierarchical Navigable Small World Graphs

**DOI:** 10.1101/2022.10.21.513218

**Authors:** Jianshu Zhao, Jean Pierre Both, Luis M. Rodriguez-R, Konstantinos T. Konstantinidis

**Author notes:** Those authors contribute equally (Jianshu Zhao, Jean Pierre Both). Corresponding author, Konstantinos T. Konstantinidis.

## Abstract

Genome search and/or classification is a key step in microbiome studies and has recently become more challenging due to the increasing number of available (reference) genomes and the fact that traditional methods do not scale well with large databases. By combining k-mer hashing-based probabilistic data structures (e.g., (Prob/Super/Densified)-MinHash or SetSketch) to estimate genomic distance, with a graph-based nearest neighbor search algorithm (called Hierarchical Navigable Small World Graphs, or HNSW), we created a new data structure and developed an associated computer program, GSearch, that is orders of magnitude faster than alternative tools while maintaining high accuracy and low memory usage. For example, GSearch can identify/classify 8,000 query genomes against all available microbial or viral genomes (n=∼318,000 or ∼3,000,000) within a few minutes on a personal laptop, using only ∼6GB of memory or less (e.g., 2.5G via SetSketch). Notably, GSearch will be even faster compared to other tools with even larger database size due to O(log(N)) time complexity and will scale well with billions of database genomes based on a database splitting strategy. Further, GSearch implements a three-step classification pipeline that accounts for the degree of novelty of query genomes relative to the database genome to maximize specificity and sensitivity. Therefore, GSearch solves a major bottleneck of microbiome studies that require genome search and/or classification of microbial or viral genomes. GSearch is available at: https://github.com/jean-pierreBoth/gsearch

## Introduction

Identifying or classifying microbial species based on either universal marker genes (e.g., 16S or 18S rRNA genes) or entire genomes represents a re-occurring task in environmental and clinical microbiome studies. However, this task is challenging because the microbial genomes in nature are still severely under-sampled by the available genomes. For instance, there are more than 10^12^ prokaryotic and fungal species in nature according to a recent estimation based on 16S rRNA gene or ITS (Internal Transcribed Spacer) analysis (1), and even more viral species (e.g. the number of viral cells outnumbers that of prokaryotic cells by a about a factor of ten in most natural habitats) (2). The number of total prokaryotic genomes (not necessarily completed, named or described biologically) has reached ∼318,000 in the newest release of the NCBI/RefSeq prokaryotic database (until Feb. 2023), and > 12 million in the latest IMG/VR4 database for viruses, representing 65,703 prokaryotic and 8.7 million viral distinct species if clustered at the 95% ANI (average nucleotide identity) level (3,4). The number of microbial database genomes is growing at a speed faster than transistors. This has created a new challenge, however; that is, identifying/searching newly sequenced genomes against these large databases to find closely related database/reference genomes for taxonomy classification has become impractical if an all-versus-all comparison strategy is applied. Further, due to the recent improvements in DNA sequencing and single-cell technologies, metagenomic surveys can now recover hundreds, if not thousands, of these yet-to-be-described genomes (or MAGs, metagenomic assembled genomes) from environmental or clinical samples in a single study (5,6), filling in the gap in the described diversity mentioned above but also further exacerbating the database size problem. In addition to the searching strategy, the actual algorithm used to determine overall genetic relatedness (e.g., ANI or its approximate) between the query and the databased genomes is critical. While the traditional Blastn-based ANI among closely related genomes at the species level, and the genome-aggregate average amino acid identity (AAI) for genomes related at the genus level or above, have been proven to be highly accurate for genetic relatedness estimation across microbial and viral genomes (7–10), they are too slow to use when dealing with more than a few dozen of genomes. Recently, phylogenetic replacements methods using a handful of universal genes (n ≈ 100) have become popular, but these methods can be memory demanding and slow (11,12), especially for a large number of or a few deep-branching (novel) query genomes. This approach cannot be applied to viral genomes, which lack universal genes. Further, universal genes due to their essentiality, are typically under stronger purifying selection and thus, evolve slower than the genome average. This property makes universal genes appropriate for comparisons among distantly related genomes, e.g., to classify genomes belonging to a new class or a new phylum, but not the species and genus levels (11,13).

Faster and more memory efficient ANI estimation based on k-mer hashing and evolutionary models have been recently described in tools like FastANI, Mash, Dashing and BinDash (14–18) that aim to alleviate this computational bottleneck. The above-mentioned tools typically rely on probabilistic data structures (or sketching algorithms) such as MinHash (a class of locality sensitive hashing) (19), HyperLogLog (20) or a combination of two, called SetSketch (21) to estimate genomic distance. Importantly, MinHash-like algorithms have been shown to be an unbiased estimation of the Jaccard index 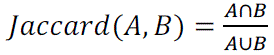 (note that Jaccard index considers only element/k-mer presence/absence in a set, unweighted) between two genomes, an accurate proxy to ANI or mutation rate after appropriate transformation (e.g., *ANI*/(1 − *Mash*) = 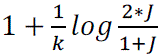, where *J* is the Jaccard index and k is k-mer size, also called Mash equation) with Poisson distribution assumption and an evolutionary model (14). However, MinHash-like algorithm can be less space efficient (e.g., use more memory) for large number of genomes than HyerLogLog, HyerLogLog++ or SetSketch despite being more accurate (21). The latter algorithms can also be used to estimate Jaccard index after estimating the set cardinality (Supplementary Note 1.) (16). However, Jaccard index (simple, unweighted) estimated by MinHash-like sketching algorithms can be problematic for incomplete genomes due to bottom-k sketch implementation (22) (14), which are commonly recovered from metagenomic surveys. Further, unweighted Jaccard estimate could be similarly inaccurate for genomes with extensive repeats such as several microbial eukaryotic genomes because this estimate does not take into account the abundance of k-mers (k-mer multiplicity). Weighted Jaccard index such as those provided by ICWS and ProbMinHash have been recently developed to address this limitation (23,24) (see also Supplementary Note 1). The application of weighted (or unweighted) MinHash-like algorithms for genome search (that is to search for the closest genomes) problem can still be slow despite the algorithms themselves being fast, because these algorithms are typically applied in a “brute force” manner. That is, all query genomes are searched against all database genomes (i.e., “all vs. all”), and thus computational time grows in a linear fashion as the number of database genome increases. More importantly, in the case of searching a database, locality sensitive hashing property of those probabilistic data structures should be satisfied to ensure high recall or accuracy.

One of the most broadly used approaches for finding closely related information to a query, while circumventing an all vs. all search, is the K-Nearest Neighbor Search (K-NNS). The K-NNS approach has been used –for instance-for 16S rRNA gene-based classification followed by a vote strategy (25). Approximate nearest neighbor search (ANNS) algorithms, such as locality-sensitive hashing (LSH) (26), k-dimension tree (27), random projection tree (28), k-graph (29) and proximity graph (30–32) have been recently used to greatly accelerate search processes with only small loss in accuracy. Proximity graph, as implemented for example in the hierarchical navigable small world graph (HNSW), has been shown to be one of the fastest ANN search algorithms(33,34) with search time complexity *O*(*log*(*N*)). HNSW incrementally builds a multi-layer structure consisting of a hierarchical set of proximity graphs (layers) for nested subsets of the stored elements. Then, through smart neighbor selection heuristics, inserting and searching the query elements in the proximity graphs can be very fast while preserving high accuracy, even for highly clustered data (31). Therefore, finding the closest genomes in a database can be substantially accelerated by combining two sub-linear algorithms while maintaining ANI/AAI accuracy: MinHash-like or HyperLogLog sketching algorithms for genomic distance estimation and HNSW for finding nearest neighbors. This is an idea that, to the best of our knowledge, has never been applied to genome search despite its potential to greatly accelerate genome search.

Here, we describe GSearch (for <u>G</u>enome <u>Search</u>), a computer program that combines one of the most efficient nearest neighbor search approaches (HNSW) with MinHash-based or SetSketch-based universal approaches to measure genetic relatedness among any microbial genomes, including fungal and viral genomes. Four MinHash-like algorithms, Densified MinHash (35,36), ProbMinHash (24), SuperMinHash (37) and SetSketch (21), due to their locality sensitive hashing property for nearest neighbor search, are provided as part of GSearch for the advantages that each provides in terms of accuracy or speed. Densified MinHash, which borrows hash values from non-empty bins to fill empty bins in the sketch vectors, which are generate based on One Permutation MinHash, is by far the fastest MinHash-like algorithm due to just one hash function used, also can be as accurate as classic MinHash. SuperMinHash is similar to classic MinHash but optimized in terms of accuracy to calculate simple Jaccard index, sacrificing speed. ProbMinHash (default) is based on shared k-mers, weighted by their abundance, and normalized by total k-mer count. Accordingly, ProbMinHash can account for genome incompleteness of prokaryotic genomes and repeats (k-mer multiplicity, or k-mer weight) commonly found in eukaryotic and sometimes in prokaryotic genomes. Essentially, ProbMinHash computes the normalized weighted Jaccard-like index *J*_+_(Supplemenatary Note 1) between each pair of genomes. SetSketch is a new data structure aiming at both space efficiency and speed, which fills the gap between MinHash (speed and accuracy, locality sensitive hashing) and HyperLogLog (space). SetSketch can be used for both cardinality estimation and set similarity (1-Jaccard index) and is also a locality sensitive hashing algorithm, providing an advantage over HyperLogLog and MinHash sketching algorithms (e.g., low memory usage and also high accuracy). The Jaccard similarity (1-Jaccard index) calculated by either of the above methods is used as input to build HNSW graph of the database genomes. Accordingly, the search of the query genome(s) against the graph database to find the nearest neighbors for classification purposes becomes an ultra-fast step and can be universally applied to all microbial genomes. The novelty of GSearch also includes a hierarchical pipeline that involves both nucleotide-level (when query genomes have close relatives at the species level) and amino-acid-level searching (when query genomes represent novel species), which provides robust classification for query genomes regardless of their degree of novelty relative to the database genomes, as well as a database-splitting strategy that allows GSearch to scale up well to billions of database genome sequences. GSearch was implemented in the Rust programming language for high performance.

## Materials and Methods

GSearch is composed of the following steps. Initially, the genetic relatedness among a collection of database genomes is determined based on the sketching algorithm Densifed MinHash, ProbMinHash, SuperMinHash or SetSketch (24,35–37), which compute the normalized weighted Jaccard-like index J_p_ using the probminhash3a algorithm (ProbMinHash option) or the simple Jaccard index J in the SetSketch/SuperMinHash/Densified MinHash options, as implemented in the probminhash package. The normalized weighted Jaccard-like similarity (1-J_p_) or Jaccard similarity (1-J) is then used as input for building HNSW graphs (note that a distance computation is required only when that genome pair is required for graph building, thus GSearch avoids all vs. all distance computations) (Figure 1a). Genomes are subsequently recursively added as the nearest neighbors of each node in the built graph file with the same distance computation procedure (Figure 1b). The built graph database and sketches from the MinHash-like algorithm are stored on disk, including a graph file from HNSW as the main component of this new data structure. Each node/genome in the graph has a corresponding sketching in the sketching file (Figure 1c). Query genomes are then searched against graph database after loading the database files and subsequently, best neighbors are returned for classification/identification. In this process, the best neighbor (or neighbors depending on the number of neighbors requested) is also identified based on the smallest ProbMinHash distance (1-J_p_) or Jaccard distance (1-J) obtained. The output can then be transformed into ANI/AAI values. The details of overall Rust parallel/concurrent implementation can be found in Figure S9. Parallel efficiency and memory usage of database building and searching can be found in Figures S10 and S11.

**Figure 1.**
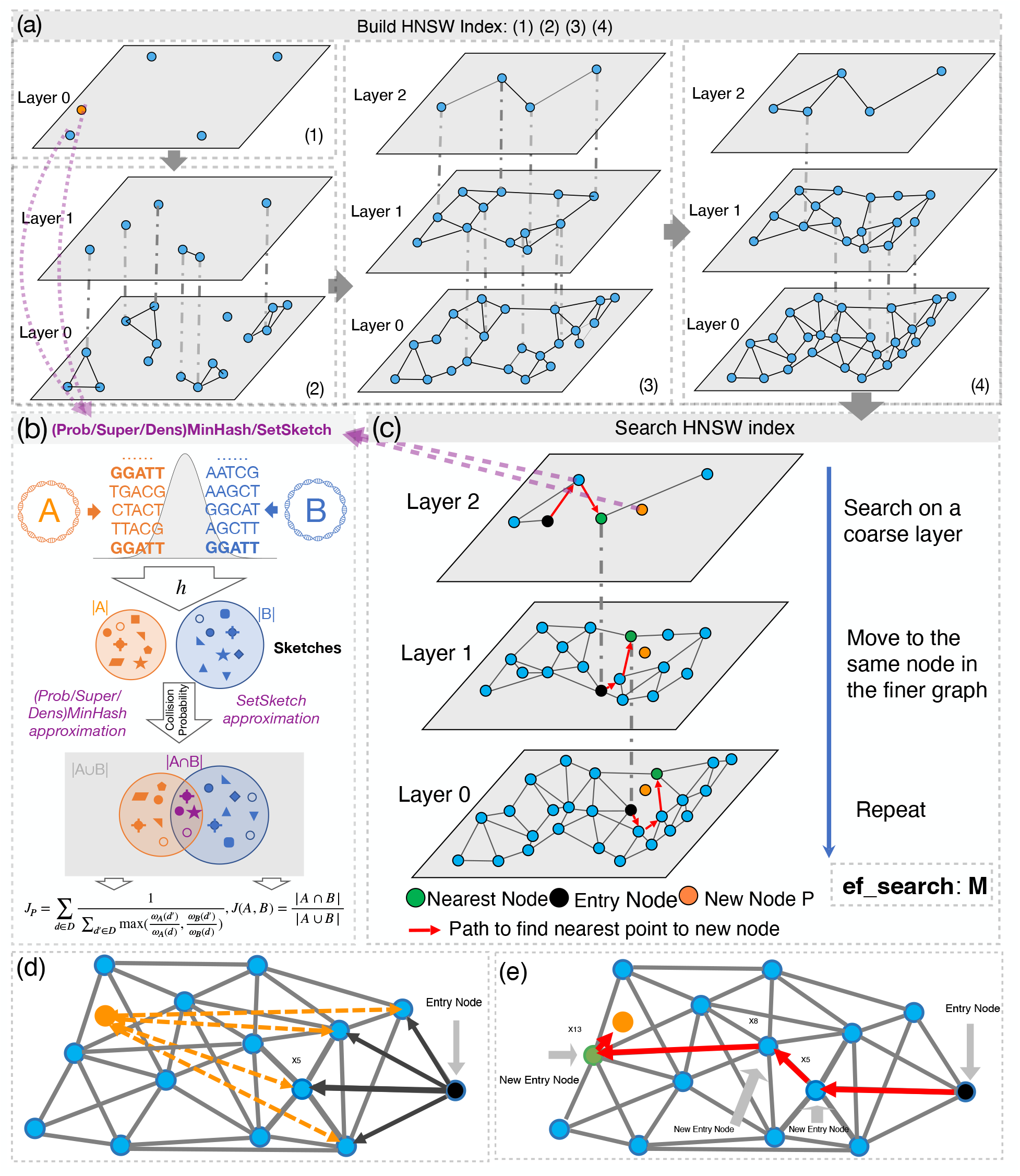
Schematic overview of GSearch building graph (a), graph searching (c) and (Super/Prob/Dens)MinHash/SetSketch distance estimated (b). (**a**) Overview of the HNSW building process. GSearch (tohnsw module) starts from a randomly chosen genome to gradually build the graph at Layer 0 (1) and gradually collapse genomes (dash grey line indicate newly clustered/collapsed representatives compare to previous step) into Layer 1 (2) and layer 2 (3) as more genomes are being incorporated in Layer 0 until all genomes in database are inserted into layer 0 (4). When inserting new genomes in the graph (database), GSearch essentially searches in a partially built graph to find nearest genomes, based on ProbMinHash/SetSketch distance values among the genomes; see (c) for details. After the required number of nearest neighbors are found for each inserted genome, a reverse update step is performed to update neighbor list of all nodes in the graph. (**b**) Overview of (Super/Prob)MinHash/SetSketch algorithms to calculate *Jp* and J, where h is the hash function to hash k-mers from two genomes (orange and blue) and store the hashed values as sketches (shapes in the two big circles). ProbMinHash calculates *Jp* from shared sketches while SuperMinHash, Densfified MinHash and SetSketch calculate J from shared sketches. (**c**) To search/identify a new genome P (orange) against the graph (request module), starting from an entry node (black, random or inherited from layer above it, depending on whether it is the top layer or not), GSearch finds the closest connected neighbor of the entry node (black) to the new node P to be searched (orange) by calculating the (Super/Prob/Dens)MinHash/SetSketch distance of the new node P (orange) with all neighbors (blue) of entry node (black) and assigns the closes one (X5 in this case) as the new entry point (that is X5 will be the new entry node) (panel (**d**)). GSearch is then traverses in a greedy manner (i.e., update the entry point using the newly found closest connected neighbor of X5) until the nearest neighbors in the layer are found (specifically, the path goes from entry node to X5, X8 and X13) (panel (**e**)), and then goes to next layer. This process is repeated until the required number of nearest neighbors (N) are all found for the given new querying data point P and subsequently, reported to the user.

### MinHash-like algorithm Implementation and Benchmark

A detailed description of the differences between ProbMinHash and traditional MinHash can be found in Supplenmentary Materials and Methods. We reimplemented the ProbMinHash algorithm in Rust to estimate genomic relatedness between any two genomes based on normalized weighted Jaccard-like similarity 1-*J_p_* according to the original ProbMinHash paper (24)(Supplementary Note 1). The MSE (Mean Standard Error) of ProbMinHash is J_p_(1-J_p_)/m, where m is the sketch size or number of registers, similar to that of classic Jaccard index J(1-J)/m. Essentially, when objects/k-mers are hashed, they represent probability distributions (Relative Frequency of k-mers after normalization), and *J_p_* is a natural extension of *J* with good properties for estimating distance among various genomes. The Rust reimplementation of ProbMinHash, and other related MinHash-like algorithms, can be found at: https://github.com/jean-pierreBoth/probminhash. We relied on version 0.1.10 of probminhash package for this study. There are 11 different MinHash-like algorithms in this library (all are metric since J and J_p_ is metric): One Permutation MinHash with Optimal Densification, Faster Densification, SuperMinHash, ProbMinHash1, ProbMinHash1a, ProbMinHash2, ProbMinHash3, ProbMinHash3a, ProbMinHash4, OrderMinHash and SetSketch (locality sensitive hashing estimator (LSH) and Joint Maximum Likelihood Estimator, JMLE). Details of the locality sensitive hashing and JMLE implementation for SetSketch, densified MinHash can be found in Supplementary Methods and Materials. Specifically, when Jaccard index is smaller than 0.01 for queries with their best neighbors/genomes found, we then use JMLE instead of LSH estimator in SetSketch since JMLE is more accurate for small Jaccard.

To benchmark ProbMinHash against Mash and Dashing 1/2 et.al., all tools were run with the same sketch size (s=12000) and k-mer size (k=16) for bacterial genomes at the nucleotide level and k-mer size (k=7) at the amino acid level for both database building and searching. For fungal genomes a larger sketch size (s=48000) was used due to much larger fungal genome sizes. Further details on choosing the k-mer size can be found in Supplementary Note 2. For convenient comparison of GSearch results against those of the Mash and ANI based methods, we performed the same transformation of Mash distance from normalized weighted Jaccard like distance J_p_ to probMASH distance as a proxy of ANI using the equation. Details of the mentioned benchmark softwares can be found in Supplementary file 3.

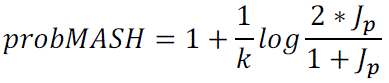

### Hierarchical Navigable Small World Graphs (HNSW)

Generally, the framework of graph-based ANN search algorithm (here HNSW) can be summarized in the following two steps: 1) build a proximity graph (HNSW) where each node represents a database vector (or genome profiles in our case). Each database vector will connect with a few of its neighbors while maintaining small world property in each layer of HNSW. 2) Given a query vector (or sequence, kmer profile in our case), perform a greedy search on the proximity graph by comparing the query vector with database vectors under the searching measures (e.g., cosine similarity or L2 similarity; in our case ProbMinHash distance). Then, the most similar candidates are returned as outputs. The key task for these two-step methods is to construct a high-quality index graph, which provides a proper balance between the searching efficiency and effectiveness. To guarantee the searching efficiency, the degree (number of maximum allowed neighbors, denoted as M) of each node is usually restricted to a small number (normally 20∼200) while width of search for neighbor during inserting (denoted as ef_construct) is usually a larger number (higher than 1000) to increase the chance to find best M neighbors by increasing the diversity of neighbors due to the large number of neighbors retained. Graph is subsequently clasped into hierarchical layers following exponential decay probability. Building graph (e.g., insert into graph) and searching query against the graph follow the same greedy search procedure except that there is an extra reverse updating of neighbors list for each vector when inserting database vectors (genomes), one by one, into the existing graph until all genomes are inserted (Figure 1a). For building, after searches are finished at the bottom layer for each inserted element, a reverse update step will be performed to update the neighbor list of each node in the existing graph, while for searching/quering against the graph this step is not needed. The overall database building time complexity is O(N*log(N)), where N is the number of nodes in the graph. The first phase of the searching process starts from the top layer by greedily traversing the graph to find maximum M closest neighbors to the new node element P in the layer by doing *ef*_construct times search (Figure 1c). After that, the algorithm continues the search from the next layer using the closest neighbor found from the previous layer as entry point, and the process repeats until to the bottom layer. Closest neighbors at each layer are found by a greedy and heuristic search algorithm (Figure 1d and e). For searching, since there is no need to reverse update best neighbor list for each node in the graph, time complexity is (only) O(log(N)) as explained above (See detailed complexity analysis on Supplementary Note 3). Theoretical guarantee of graph-based algorithm can be found in Supplementary Note 5. We reimplemented the original hnswlib library written in C++ using the Rust programming language for memory safety, efficient parallelism and speed. We also implemented memory map in the newest version, a feature not in the original C++ library that is useful for billions of data points or genomes. The implementation can be found at https://github.com/jean-pierreBoth/hnswlib-rs. Version 0.1.19 was used in this study. Benchmarking of this package against standard datasets and comparison with other implementations can be found in the Supplementary Materials and Methods.

### Details of program implementation in Rust

There are 3 modules in total: tohnsw, add and request. Tohnsw is to build the graph by gradually inserting genomes into graph while request is to query new genomes against the graph database built in the tohnsw step. Tohnsw starts from reading database genomes and generating k-mer profile and sketches using ProbMinHash3a algorithm (or SuperMinHash, SetSketch and Densified MinHash) for distance calculation (Figure 1a and c). Next, tohnsw selects, at random, the first batch of genomes to insert into the graph (Figure 1a (1)), following HNSW constructing rules mentioned above and taking into account the computed ProbMinHash distance (1-J_p_) or SetSketch/SuperMinHash/Densifed MinHash approximated Jaccard distances (1-J) between genomes to connect genomes based on their relatedness (Figure 1b) until all genomes in database have been inserted (Figure 1a (2), (3) and (4)). Finding nearest genomes for the genome to be inserted is essentially a search process but search in a partially built graph, which is similar to the request/search module: whenever a genome is going to be searched against the existing graph, each genome in the graph is associated with a list that stores the M closest neighbors/genomes to the genome and the distance to these neighbors. Then, the distances of this genome with the nearest neighbors (M) of entry genome in each layer will be computed (ef_construct times) using probminhash3a algorithm or SetSketch and the smallest distance of the neighbor genomes will be the new entry genome (Figure 1d and e). This process will be repeated until the nearest genomes (<=M) in the layer are found and subsequently, the program will go to the layer below, using the genome that was represented by the nearest genome in the above layer as new entry genome in the new layer. The search layer algorithm is repeated until the bottom layer is reached/analyzed (Figure 1c). In contrast to the default settings in the original hnswlib, we allow the two parameters of neighbor selecting heuristics, *extendCandidates* to be true and *keepPrunedConnections* to be false because our genomic data is extremely clustered and there is no need to fix the number of connections per element considering the maximum connection allowed. The “Add” module is to add new genomes to pre-built HNSW database using default parameters loaded in the pre-built database files. Request module will load the graph database and then search query genomes against it to return the best neighbors of each query, following exact the same procedure with building step without updating the database. Tohnsw, add, and request modules are all operating in parallel for high performance (see Supplementary Note 6 and Table S9, S10 and S11). GSearch relies on kmerutils v0.0.10 (https://github.com/jean-pierreBoth/kmerutils), which is a Rust package we developed to manipulate genomic fasta files including k-mer string compression, recursive k-mer hashing, k-mer counting, filtering using cuckoo filter. All tests performed in this study were based on GSearch version 0.1.3. Scripts for reproducing the results of this study can be found here: https://github.com/jianshu93/gsearch_analysis. The GSearch package can be found here: https://github.com/jean-pierreBoth/gsearch.

### Prokaryotic classification pipeline

The amino-acid level graph showed that the closest database neighbors were found, with high recall, when the query shared at least 52% AAI to its best neighbor. For more divergent genomes (<52% AAI), whole-genome amino-acid level graph loses accuracy and we had to switch to universal, single-copy protein-coding genes. For the nucleotide-level graph, we used k-mer=16 for bacteria and archaea to have high specificity for closely related database genomes (e.g., sharing about 95% ANI). For building the whole-genome amino-acid graph, we used k=7 to have the best specificity without compromising sensitivity, which is also consistent with previous results on classification of amino acid sequences based on k-mers (38). For building graph based on universal gene set, we use k=5 because of much smaller total amino acid size. For further details on the range of k-mer to use for bacteria genome and proteome, and viral genome and proteome, see Supplemental Notes 2.

The proteome of each genome was predicted by FragGeneScanRs v0.0.1 for performance purposes as opposed to Prodigal software despite small loss in precision (39) (Supplementary Table S11). Hmmsearch in the hmmer (v3.3.2)(40) software was used to extract the universal gene set for bacteria and archaea genomes (universal gene graph). Note that for viral genomes, this last step was not implemented because there are no universal single copy genes for viral genomes. Evaluation of the speed and memory requirements for all steps mentioned above were performed on a RHEL (Red Hat Enterprise Linux) v7.9 with 2.70 GHz Intel(R) Xeon(R) Gold 6226 CPU. Unless noted otherwise, all 24 threads of the node were available by default.

### Distributed implementation and database splitting

To accommodate the increasing number of genomes that become available at an unprecedented speed in recent years and will soon reach 1 million or more microbial genomes, we provide an option to randomly split the database into a given number of pieces and build graph database separately for each piece. In the end, all best neighbors returned from each piece will be pooled and sorted by distance to have a new best K neighbor collection returned to the user for each query genome. We hereby prove that in terms of requesting top K best neighbors, the database split strategy is equivalent to non-split database strategy if the requested best neighbors for each database piece is larger than or equals to requested best neighbors in the non-splitting strategy. The underlying reason is that the best neighbors for a given query globally (non-splitting) are also the best locally for the same piece of database (e.g., best 5 neighbors in piece #1 locally –meaning only in this piece-maybe also be the best n=<5 neighbors genomes globally). The database splitting and request can be done sequentially, on a single node, without multi-node support. In theory, a large database can be split into any number of pieces. In practice, a reasonable way to decide on the number of database pieces to use is so that memory requirement for each piece is equal or smaller than the total memory of host machine.

### Species database and testing genomes for benchmarking and recall

GTDB version 207 and entire NCBI/RefSeq prokaryotic genomes (as of Feb. 2023) was used to build the database for bacteria and archaea genomes. The IMGVR database version 4, with species representatives at a ≥95% ANI, was used to build the database for viruses (a total of ∼3 million). For fungal genomes, all genomes downloaded from the MycoCosm project (as of Jan. 2022) were used (41). The amino acid sequences of predicted genes on the genomes were obtained using FragGeneScanRs for bacteria/archaea and GeneMark-ES version 2 for fungi (42). The Universal Single Copy Gene (USCG) gene set for GTDB genomes were extracted via hmmer software (40).

To test the performance of our pipeline, we specifically chose genomes that are not included in the GTDB database (the database was used for graph building). In particular, the bacterial/archaeal genomes, mostly MAGs, reported by Ye and colleagues (43) and Tara Ocean MAGs (total 8,466 MAGs) (44) were used. We randomly selected 1000 genomes/MAGs from Ye’s collection and use them as query genomes to test the performance and accuracy of GSearch.

### Recall of AAI-, ANI– and MinHash-based nearest neighbor searching for bacteria/archaea, fungi and viral genomes

To benchmark GSearch performance compared to traditional ANI/AAI– and/or MinHash-based tools, we ran brute-force Blastn-ANI/FastANI (MUMMER for fungi ANI), Blastp-based AAI to find the best neighbors for the same query genomes in different dataset and evaluated whether the best neighbors found by GSearch were the same as those found by Blastn-ANI/FastANI and blastp-based AAI. Briefly, we used the well-known average recall as the accuracy measurement (45). Given a query genome, GSearch is expected to return k best genomes. Then we examined how many genomes in this returned set were among the true k nearest neighbors (NN) of the query found by the reference brute-force Blastn-ANI and/or Blastp-based AAI. Suppose the returned set of k genomes given a query is R’ and the true k nearest neighbors set of the query (from Blastn-ANI or blastp-based AAI) is R, the recall is defined as: 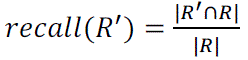. Then, the average recall represents the mean of all the query datasets/genomes. We compare the accuracies of different algorithms by requiring different number of nearest neighbors (NN) of each query genome, including 10-NN and 5-NN, using the same approach. Since biological species databases are generally sparse due to the undersampling of natural diversity and the existence of natural gaps in diversity among species, a larger top k NN (e.g., 100) used in standard ANN benchmark experiments will offer little, if any biological advantage, especially when the query genomes are relatively new, e.g., a new family compared to database genomes. Therefore, we use top 5-NN and 10-NN. Further, if the genomic distance (J_p_) of query to some of the top 10 or top 5 neighbors found by GSearch at the nucleotide level was larger than 0.9850 for bacterial genomes, these matches were filtered out and not used in estimation of recall, because we have shown that above this threshold, k-mer based MinHash methods at nucleotide level will lose accuracy and this is not related to HNSW search itself (e.g., 8 out of 10 are kept, so only top 8 in R’ is compared with top 8 in R). At proteome level, accuracy was calculated following similar rules with the threshold value being 0.9720 (switching to the universal gene graph at this level). For viral genomes, the threshold was 0.9800. Those thresholds were chosen based on correlation with blast-based ANI or AAI. Detailed commands for running each software mentioned above can be found in supplementary file 3.

## Results

### ProbMinHash as a robust metric of genome relatedness for prokaryotic genomes

Correlations between ProbMASH (we called it ProbMASH, after transformation from ProbMinHash distance (Jp), see Materials and Methods for details) and ANI (determined by FastANI) or Mash distance showed that ProbMinHash is robust and slightly better than Mash for determining distances among bacterial genomes related at ∼78% ANI, or above, i.e., genomes assigned to the same or closely-related species (Spearman rho=0.9643 and 0.9640 for ProbMinHash and Mash values against the corresponding ANI values for the same genome comparisons, respectively, *P*<0.001, Figure S1a and S1b). For moderately related genomes, for which ANI based on nucleotide k-mer is known to lose accuracy, ProbMASH (ProbMinHash) based on amino acid k-mer was robust compared to Blast-based average amino acid identity or AAI, especially among genomes showing between ∼52% and 95% AAI (Spearman rho=0.90, *P*<0.01, Supplementary Figure S2a and S2b). Below ∼52% AAI, both ProbMinHash and Mash distance lose accuracy compared to AAI. However, AAI of just universal genes provides a robust measurement of genetic relatedness at this level of distantly related genomes, and we show here that ProbMASH (from ProbMinHash distance) for this set of universal genes is also robust (Spearman rho=0.9390, *P*<0.001, Supplementary Figure S3). Thus, for query genomes of organisms with only distant relatives in the database (i.e., deep branching), for which their closest matching genome in the database is related at the order level or higher, restricting the search to the universal genes can provide robust classifications (11). Accordingly, GSearch implements a three-step classification process, depending on the degree of novelty of the query genome against the database genomes, using the AAI thresholds mentioned above (see also Fig. 2g). This strategy and its accuracy are discussed further below. To show the accuracy of One Permutation MinHash with Optimal Densification (Densifed MinHash), we calculated ANI by both Mash (classic MinHash) and our implementation, we showed that they are almost identical (Figure S12).

**Figure 2.**
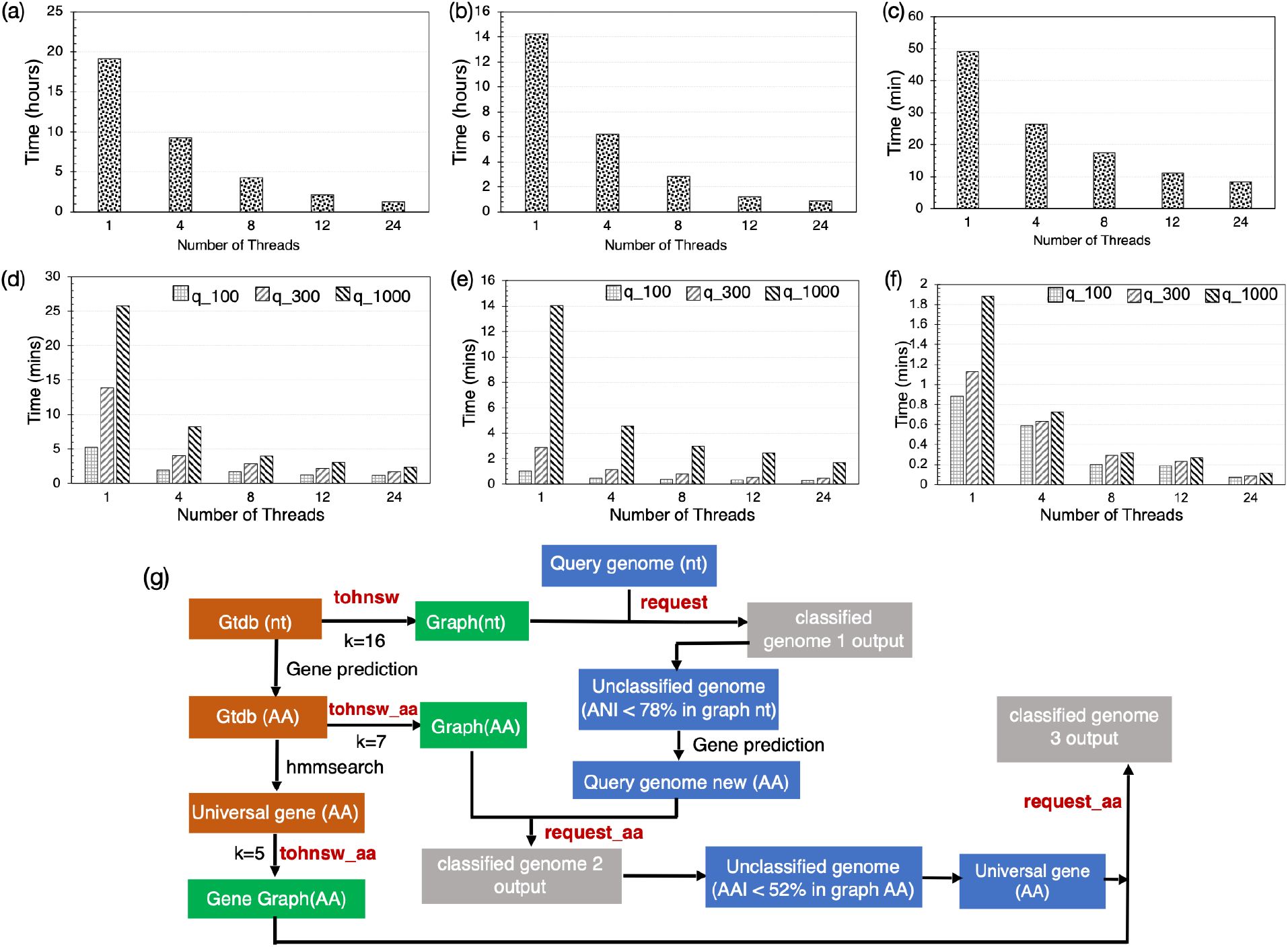
Performance and scalability of database building (tohnsw) and searching (request) steps as the number of threads increases. Upper panel shows total wall time (y axes) for building the GTDB v207 (65,703 genomes) at the nucleotide level **(a)**, whole-genome proteome (amino acid level) **(b)** and universal gene set proteome **(c)** reference graphs (or databases). Middle panel shows searching performance and scalability against number of threads used for different sets of query genomes at nucleotide **(d)**, amino acid level **(e)** and universal protein level **(f)**. Specifically, 100, 300 and 1000 query genomes (figure key) were used. All tests were run on a 24-thread Intel (R) Xeon (R) Gold 6226 processor, with 40GB memory available. **(g)** Overview of GSearch’s the 3-step pipeline for classifying prokaryotic genomes. Orange boxes denote steps that aim to prepare genome files, in different formats, for graph building while green boxes denote building steps of the graph database (in nucleotide or amino acid format). Blue boxes indicate input/query genomes to search against the database while grey boxes indicate classification output for each input. Two key steps of GSearch: tohnsw and request are used to build graph database and request (or search) new genomes against the database, respectively. Two thresholds are used in the pipeline to decide between whole nucleotide vs. whole-genome amino acid search and whole-genome amino acid vs. universal gene amino acid; that is, 78% ANI and 52% AAI, corresponding to Probminhash distance 0.9850 and 0.9375, respectively (see main text for details).

### Comparisons of sequence search algorithms and MinHash-like algorithms

We compared GSearch with other general-purpose sequence or genome search algorithms, focusing on time complexity (big O annotation) of each sequence or genome search algorithm. GSearch was, by far, the fastest genome search algorithm, with a O(log(N)) complexity (Table 1). Other general-purpose sequence search algorithms (e.g. Sequence Bloom Tree, COBS et.al.) are either sublinear under certain assumptions or not practical in real world applications. Also, none of the general-purpose sequence or genome search algorithms were previously benchmarked against an ANI-like distance; hence, their accuracy remains untested.

**Table 1.** Bioinformatics algorithms for sequence/genome searching and their comparisons. S represents a document/genome. N is the total number of documents/genomes. U_s∈S_ |s| represents the total number of terms in N documents/genomes and ∑_s∈S_ |s| is total unique terms/k-mer. MPH is minimal Perfect Hashing. For the Inverted Index size, the extra log (N) comes from the bit precision document IDs. For SBTs, log(N) is the height of the tree, and for Bloom Filters at each level is *O*(∑_s∈S_ |s|) in total. MinHash time and space complexity was based on k hash functions for N sets, each set has d’ non-zero element. Note that the most recent B-bit MinHash with optimal densification can further improve to O(N*(d’+k)). O(vdlog(N)) for GSearch is O(log(N)) in practice since v and d is small and single pair genome comparison is a constant (Supplementary Note 3). τ depends on the user-specified parameter and is generally less than 0.01 in practice.

|  | Query Time | Space | Recall Benchmark (Biologically) | Comments |
| --- | --- | --- | --- | --- |
| Inverted Index<sup>1</sup> | $O(1)$ | $Best: \log(N) \cup \vert s\vert $ | Only for document search/retrieval, could be applied to genomic/sequence search | **Long construction time; impractical for bigger datasets; best case needs MPH and a known k-mer (term) distribution** |
| BIGSCI/COBS<sup>2,3</sup> | $O(N)$ | $\sum_{S \in S} \vert s\vert $ | A hybrid between an inverted index and Bloom filters (COBS), high false positive rate, no benchmark with mutation rate by ANI/Mash | Query time is linear in $N$, small index size |
| Sequence Bloom Tree<sup>4</sup> | $Best: O(\log(N)),$<br>$Worse: O(N)$ | $\log(N) \sum_{S \in S} \vert s\vert $ | Given the k-mers from a query sequence, the task is to determine which of the $N$ documents contain all the k-mers present in the query; no benchmark with mutation rate by ANI/Mash | **Sequential query process is bottleneck; designed for sequential implementation** |
| RAMBO<sup>5</sup> | $O(\sqrt{N} * \log(N))$ | $\Gamma \log(N) \sum_{S \in S} \vert s\vert $ | Similar to SBT, finding which of the $N$ documents/genomes contain all the k-mers present in the query, no benchmark with ANI/Mash | Only for $\Gamma < 1$, query time is sub-linear |
| MinHash<sup>6</sup> | $O(N * d' * k)$ | $O(N * k)$ | Average Nucleotide Identity (ANI) or mutation rate via Mash distance | Query time is linear in $N$ |
| GSearch (ProbMinHash+HNSW) | $O(vd\log(N))$ | $O((1 + \tau) * N * d + k * N)$ | Average Nucleotide Identity (ANI) via Mash-like mutation rate/index | Long database construction time $O(N * \log(N))$, but users are free from construction. |
| FLINNG<sup>7</sup> | $O(\log^2(\frac{1}{\delta}) \log^3(N) N^{\frac{1}{2}+\gamma})$ | $N^{\frac{3}{2}} \log^2(N)$ | No Benchmark with Average Nucleotide Identity (ANI) via Mash-like mutation rate/index, only 15% of RefSeq genome meet the γ-stable criteria | The γ-stable query condition is a relatively strong requirement for the query. Limitation: works for queries for which the neighbors are all above a (relatively high) similarity threshold to the query |

<sup>1</sup>Croft, Metzler and Strothman <sup>1</sup>, <sup>2</sup>Bradley, Den Bakker, Rocha, McVean and Iqbal <sup>2</sup>, <sup>3</sup>Bingmann, Bradley, Gauger and Iqbal <sup>3</sup>, <sup>4</sup>Solomon and Kingsford <sup>4</sup>, <sup>5</sup>Gupta, Yan, Coleman, Kille, Elworth, Medini, Treangen and Shrivastava <sup>5</sup>, <sup>6</sup>Broder <sup>6</sup>, <sup>7</sup>Engels, Coleman and Shrivastava <sup>7</sup>

We then also compared available MinHash-like sketching algorithms in terms of their time complexity (big Oh), space (memory), variance, and other important properties for large scale applications such as mergeability (whether sketching part of dataset and then merge is equivalent to sketching the entire dataset) to estimate Jaccard-like index (Table S1 and S2). SuperMinHash and ProbMinHash are fast and mergeable MinHash algorithms, slower only than One Permutation MinHash with Optimal Densification (as implemented in BinDash) (Table S1). The latter algorithm, however, is not mergeable thus difficult to apply in a distributed computational environment. HyperLogLog-like algorithms are significantly more space efficient albeit slightly slower. For variance, we analyzed the rooted mean square (RMSE) error of all MinHash-like sketching algorithms mentioned above. MinHash-like algorithms such as classic MinHash, ProbMinHash, SuperMinHash had the smallest theoretical variance as m (the number of registers used for sketching) increases (we use m=12,000, note that J*(1-J) is much smaller than 1 for all Jaccard), following by Densified MinHash, SetSketch (as b→1) (Table S2). Our implementation of the Densified MinHash has almost close RMSE with classic MinHash (Figure S13). SetSketch is also space efficient and is similar to that of HyperLogLog in terms of space and speed. HyperLogLog sketch estimators such as those implemented in Dashing has largest variance for estimating the cardinality of sets (followed by inclusion-exclusion rule to estimated Jaccard index, Supplementary Note 1), consistent with experimental results when searching query genomes against database (Supplementary Table S14 and S15), especially for small Jaccard distance (e.g., 75% to 78% ANI, corresponding to 0.009 and 0.015 Jaccard index), where variance is more important for accuracy.

### Graph building and speed of search against reference prokaryotic genomes

To build the database graph for the entire GTDB v207 database at the nucleotide level (65,703 unique, non-redundant prokaryotic genomes at the 95% ANI species level), the tohnsw module (database build subcommand of GSearch) took 1.3 h on a 24-thread computing node and scaled moderately well with increasing number of threads (0.27 h using 128 threads) (Figure 2a). Maximum memory (RAM) required for the building step was 15.3 GB. The total size of written database files on disk was ∼3.0 GB. There are 3 layers for the resulting graph, with 65180, 519, and 4 genomes for layer 0,1 and 2 respectively. The searching of query genomes against the GTDB database graph, requesting best 50 neighbors for 1000 query genomes, which represented different previously known as well as novel species of eight bacterial phyla (see Methods for details on query genome selection), took 2.3 min (database loading was 6 s) on a 24-thread machine and scaled well with increasing number of threads (Figure 2d). The memory requirement for the request (search) step was only 3.0 GB for loading the entire database file in memory.

We also built a database of all NCBI/RefSeq prokaryotic genomes (∼318K, 2 Terabytes in total), which took 4.1 hours using 24 threads with maximum memory usage of ∼21GB (1.2 h with 128 threads) and dumped database file size of 15GB (30GB for amino acid). Searching of 8,466 query genomes against this RefSeq database took 9.32 min (Table 2) and ∼16GB of memory, which was significantly better than alternative state-of-the-art tools for the same purposes such as Dashing1/2 and BinDash (brute force HyperLogLog or MinHash), which took 28/42 min and 21 min, respectively (Table S3, Figure S5). To evaluate the performance against Dashing or Sourmash more fully, we increased the number of database genomes gradually from 60K to 318K (all RefSeq prokaryotic genomes, 2 Terabytes) while using the same number of query genomes (8,466). We observed that GSearch follows a log fitting, consistent with the O(log(N)) theoretical prediction, while Dashing 1/2 and Sourmash follow linear fitting, and thus the latter tools will be increasingly slower than GSearch as the number of database genome increases (Figure 3a and c). For example, GSearch was 3 to 4 times faster than Dashing 1/2 and skani when the number of database genomes increased from 65,703 to 318,000 (Figure 3a, Table S3) and 20 to 30 times faster with the 3 million reference virus genomes (Figure 3b, Table S4. see also below). The speedup will be even larger as number of database genomes continues to grow. With Densifed MinHash as option, GSearch can be 2-3 times faster than ProbMinHash (Table S3)

**Figure 3.**
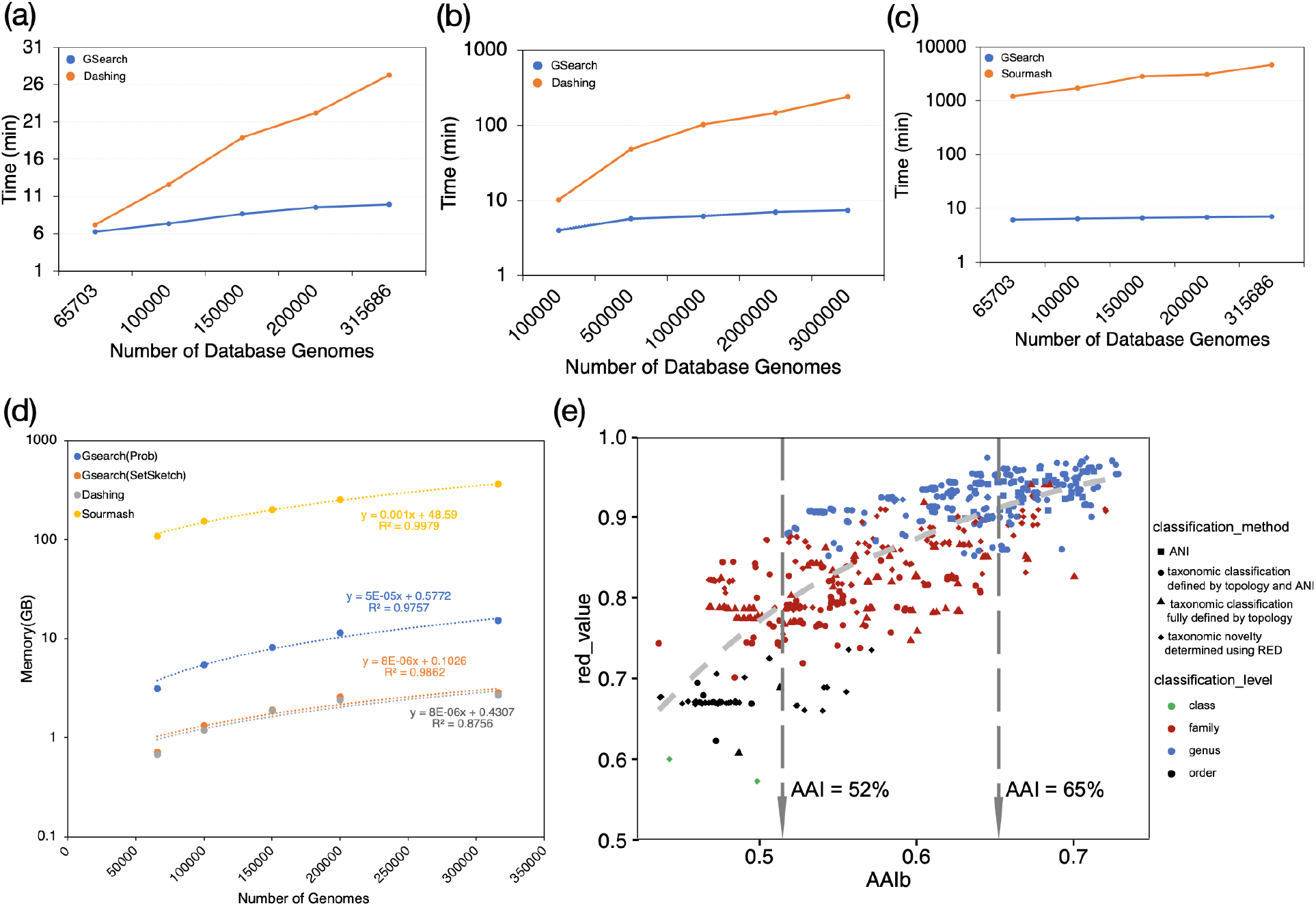
Running time, memory consumption and classification accuracy of GSearch against Dashing, Sourmash and Blast-based ANI/AAI tools. (**a**) Running time of GSearch versus Dashing for searching 8,466 query genomes against the RefSeq prokaryotic genome database as a function of the number of genomes used in the database (x-axis) at the nucleotide level. **(b)** Running time of GSearch (blue) versus Dashing (orange) for 10,000 query viral genomes against the IMG/VR v4 database at amino acid level. **(c)** Same as in (a) above but comparison is against Sourmash (orange). **(d)** Memory consumption of GSearch during versus Dashing and Sourmash search. Search is to load database into memory, thus maximum memory is directly related to database size. Since Sourmash search is not parallelized, GNU parallel was used for process-level parallelism. **(e)** Comparison of GSearch classification results with GTDB-Tk and Blastp-based AAI tools for moderate-to-distantly related query genomes based on the bacterial proteome database (e.g., ANI between the query genome and its best match in the database was lower than 78% for these genomes). Each point represents a comparison between two genomes, query and the best match found by GSearch, showing RED values generated by GTDB-tk (y-axis) versus Blastp-based AAI between these two genomes. The taxonomic rank that the query and the best database match share is shown (see figure key). Two vertical lines indicate Blastp-based AAI threshold for family and genus level classification threshold. Note that the best match was always the same genome between GSearch and all vs. all Blastp AAI, and the overall consistency between GSearch/AAI and GTDB-tk in identifying the same best database genomes for the same query genomes. Note that y-axis values are in log scale in panels b, c, and d.

**Table 2.** GSearch’s performance on major CPU platforms for searching 8,466 query genomes against the RefSeq reference database (∼318K genomes). ProbMinHash was used as the genomic distance algorithm for this benchmark, with 50 neighbors requested.

| CPU | Number of threads | Clock speed (GHz) | Request time for nt (min) | Gene Prediction-FGSrs (min)<sup>c</sup> | Request time for proteome (min) | hmmsearch time (min)<sup>d</sup> | Request time for USCG (min)<sup>d</sup> |
| --- | --- | --- | --- | --- | --- | --- | --- |
| Intel (R) Xeon (R) Gold 6226<sup>a</sup> | 24 | 2.70 | **9.329** | 3.348 | **6.234** | 3.524 | **0.445** |
| Intel (R) Core i9-8950HK<sup>b</sup> | 16 | 2.30 | **13.654** | 6.764 | **7.91** | 6.214 | **0.613** |
| AMD EPYC 7513a<sup>a</sup> | 32(24 used) | 2.60 | **10.245** | 4.452 | **6.021** | 3.345 | **0.402** |
| Apple M1 Pro<sup>b</sup> | 10 | 3.22 | **9.669** | 4.31 | **6.166** | 4.603 | **0.568** |

<sup>a</sup>RHEL v7.9, Linux v3.10.0-1160, all threads used. 512G RAM available.
<sup>b</sup>MacOS v12.3, Darwin 21.4.0 on MacBook Pro laptops, all threads used. Database was split into two pieces to have smaller memory requirement on those 16GB RAM laptops since database size was ~15GB for entire NCBI/RefSeq.
<sup>c</sup>Parallel package was used to run multiprocess at the same time. FGSrs stands for FragGeneScanRs (comparisons with Prodigal can be found in Table S11). Note that in practice only those genomes failing in the request step at the nucleotide (nt) level (best match found was less than 78% ANI) will be used in this amino-acid step.
<sup>d</sup>Only 1000 genomes were used for testing hmmsearch and USCG request in GSearch because this step was for assessing a few genomes that were novel at the order level or above. Parallel Packages was used to run multiple processes of hmmsearch, one thread per process for hmmsearch.

To build the amino-acid-level graph for moderately related query genomes, all GTDB v207 genomes were used for gene calling by FragGeneScanRs and subsequently, the predicted amino acid sequences for each genome were used for building the database graph. The graph building step took 1.4 h (Figure 2b) with a maximum memory required to be 37.7 GB on the 24-thread node. The total size of the written database files on disk was 5.9 GB. However, when the SetSketch option was used, graph building step took 2.3 h but only 9.4GB maximum memory needed, with the database size being 0.5GB (Figure 3 (d)). There were 65158, 543 and 2 genomes for layer 0,1 and 2 respectively. The entire RefSeq genome database (∼318K) took 3.7 hours to build at the proteome level with similar maximum memory, dumped files size was 29GB (only 2.5GB with the SetSketch option). Requesting 50 neighbors for querying 1000 genomes at the amino-acid level took 1.52 minutes with a memory requirement of ∼6.0 GB (database loading 9 seconds; Figure 2e) for the GTDB database. Requesting 50 neighbors for 8,466 query genomes against the NCBI/RefSeq proteome graph, took about 6.2 min (Table 2). In comparison, Mash dist took 207.2 min against NCBI/RefSeq with 24 threads for the same task (no amino acid option for Dashing).

Finally, for most distantly related query genomes, the graph building for the universal gene set follows the same logic as the amino acid level graph mentioned above except for using a smaller k-mer size (k=5) due to the smaller k-mer space of ∼120 universal genes vs. the whole-genome level. It took 7.76 min to build the database (Figure 2c) and 12 seconds to request 50 neighbors for 1000 queries on a 24 threads node (Figure 2f) against the GTDB database graph. Building for the entire NCBI/RefSeq database took about 27 min while requesting 1,000 against it took about 0.445 min (Table 2).

### Searching accuracy and speed for prokaryotic genomes

The accuracy of GSearch and other tools in identifying the best matching genomes among the database genomes was evaluated by comparing against the best matching genomes for the same query genome(s) identified by traditional alignment-based methods, namely Blast-based ANI for query genomes with close relatives in the databases and Blast-based AAI for query genomes distantly related to their best match in the database. K-mer-based genomic distance (e.g. all tools mentioned above) is widely known to be less accurate with higher degree of novelty of the query genomes relative to the database genomes as also explained in Supplementary Note 2 and 4. To address this limitation, GSearch employs a three-step framework, calculating genomic distance at nucleotide, whole-genome amino acid (proteome) and universal gene amino acid levels for query genomes of increasing novelty (or genetic distance) to database genomes as described above (Fig. 2g). For query genomes from two datasets, the Tara Ocean (8,466), and the collection put together by Ye and colleagues (997), that had closely related genomes in the RefSeq database (showing >78% ANI, 6,992 and 906 query genomes, respectively), we observed an average top 10 recall of 96.2% and 95.1%, respectively. Recall was estimated by comparing (meaning % overlap) GSearch’s top-10 matching genomes to top-10 found by Blastn-ANI (similar results were obtained for top-5 matching genomes; Table 3). For query genomes with no closely related genomes in the database showing higher than 78% ANI, 1474 and 91 genomes from the two datasets, respectively, recall was significantly lower, only around 60% or below (Table 3). However, when we search these genomes at the proteome level using Blastp-AAI as the reference standard for calculating recall, the recall value was 96.9% and 95.2%, respectively (Table 3). There were 327 and 25 genomes, respectively, that, even at proteome level, they had no related genome in the RefSeq database showing higher than 52% AAI and these genomes accounted for most of differences with the top-10 matching genomes by blast-AAI. These genomes apparently reflect the fact that the public genome databases are still sparse and do not cover well all biological diversity. When we searched these deep-branching (highly novel) genomes at the universal gene proteome level, using the universal gene AAI as the reference standard, we found the average top-10 recall value of 95.5% and 94.6%, respectively (Table 3). We also compared these results obtained by GSearch to those of BinDash, Dashing 1/2 and Sourmash for the same query genomes. We found that Sourmash is as accurate as GSearch but much slower, while Dashing is less accurate than GSearch and Sourmash or BinDash, especially when the closest database genomes showed less than 80% ANI to the query genomes (moderately or distantly related) (Table S14 and S15). The results were similar independent of the estimator used in Dashing (e.g., MLE or JMLE), consistent with theoretical predictions (Table S2). BinDash was as accurate as Mash but much faster as previously described (Table S1 and Table S13). We provide a theoretical derivation based on k-mer conservation and empirical results for deriving the 78% ANI and 52% AAI thresholds to decide the switching between whole-genome nucleotide vs. amino acid (proteome) search and whole-genome proteome vs. universal gene only, respectively, in Supplementary Notes 2 and 4. Detailed accuracy results for all tools based on a single query genome (not average recall across all query genomes as shown above) can be found in Supplementary Table S13, S14 and S15.

**Table 3.** Recall of GSearch for two query genome datasets containing various levels of novel genomes relative to the genomes in the reference database. The query genome datasets are the same as those used in Table 2. Blastn-ANI, blastp-AAI and universal gene blastp-ANI were used as the reference results (or standards) to compare against. Recall is the average across all query genomes.

| Dataset | Dataset size<br>(# of<br>genomes) | level | Recall (top 5) | Recall (top 10) |
| --- | --- | --- | --- | --- |
| **Tara (with best in database)<sup>1</sup>** | **6,992** | **nucleotide** | **98.3%** | **96.2%** |
| Tara (without best hit in database) | 1474 | nucleotide | 62.4% | 43.1% |
| **Tara (with best in database)<sup>2</sup>** | **1147** | **proteome** | **97.2%** | **96.9%** |
| Tara (without best hit genome in database) | 327 | proteome | 65.0% | 50.8% |
| **Tara (with best in database)<sup>3</sup>** | **327** | **universal** | **96.4%** | **95.5%** |
| **Ye (with best in database)<sup>1</sup>** | **906** | **nucleotide** | **97.7%** | **95.1%** |
| Ye (without best hit in database) | 91 | nucleotide | 54.8% | 48.5% |
| **Ye (with best in database)<sup>2</sup>** | **66** | **proteome** | **96.3%** | **95.2%** |
| Ye (without best hit in database) | 25 | proteome | 76.0% | 55.5% |
| **Ye (with best in database)<sup>3</sup>** | **25** | **universal** | **95.7%** | **94.6%** |

<sup>1</sup>Defined as query genomes that had best match in the database better than 78% ANI; <sup>2</sup>defined as query genomes that had best match in the database better than 52% AAI; <sup>3</sup>defined as query genomes that had best match in the database better than 55% AAI based on universal genes.

We also evaluated the effect of genome completeness on the accuracy of GSearch given that bacterial genomes recovered from environmental metagenomic surveys are frequently incomplete. GSearch was robust to genome incompleteness down to 50% completeness level. We found that at genome completeness higher than 50%, the accuracy/recall in the top 10 best matches was higher than 80% and decreased considerably below this completeness level (Table S12). Hence, GSearch analysis is not recommended for genome completeness less than 50% due to the normalization step in ProbMinHash.

### Graph database building and searching accuracy for viral and fungal genomes

Graph building and requesting for viral genomes is not effective at the nucleotide level because many viral genera are genetically too distant from each other and do not have close relatives in the public genomic database; that is, the database is too sparse. Accordingly, k-mer-based methods for genomic distance will often lead to imperfect graph structure for viral genomes. Therefore, we built only an amino acid level graph for viral genomes, using all genes in the genome due to the lack of universal genes for viral genomes. Database building took 13.89 h for all ∼3 million IMG/VR4 genomes on a 24-thread node and graph file on disk was 15.8 GB (Figure S6a). Requesting 1000 top neighbors scaled well with increasing number of threads and took about 3.63 min (database loading took an additional 1.1 min) using 24 threads (Figure S6b). The top-5 neighbors for 1000 query phage genomes were highly overlapping (98.32% recall; Table S6) compared with the top-5 neighbors of the brute-force AAI approach. For such large databases in terms of number of genomes, GSearch is about 20X faster than the brute-force Mash and Dashing tools in finding the top-5 neighbors (Figure 3b, Tables S4). We also compared GSearch with a new database building method, called PhageCloud, which relies on manually curated genome labels (e.g., environmental source) for graph database building using the Neo4j software and Dashing for genetic distance computation. Since PhageCloud provides only a web implementation with one genome query at a time, we limited the search to one viral query genome against the same database used by PhageCloud i.e., Gut Phage Database ^38^. It took 37 seconds to find the two best matches with PhageCloud while GSearch took 15 seconds (database loading 14 seconds, search 1.5 second) for the same search. Mash, on the other hand, took 4 minutes to find the same two best matches. It should be noted, however, that, because the database is already available (loaded) on PhageCloud’s website, 37 seconds is only for the searching step and website responses (average value for 5 runs on 5 different days) whereas GSearch took only 1.5 second for this same step.

Graph building for fungal genomes is slower compared to prokaryotic genomes, despite the smaller number of available fungal genomes (n=9700), because the average fungal genome size is much larger and k-mer and sketch size are accordingly much larger (k=21, s=48000). It took 2.3 h on a 24-thread node to build the nucleotide level graph for these fungal genomes. Searching step was also slower due to the larger k-mer space. Accordingly, it took 3.13 min to request 50 neighbors for 50 query fungal genomes while Mash tool 4.4 min. Nonetheless, top 10 recall was still very high (∼99.4%) against Mash and MUMMER-based ANI for the same datasets. For the amino acid level graph, the time for graph building was only 0.61 h, shorter than the corresponding prokaryotic graph. Identifying 50 neighbors for 50 query fungal genomes at the amino-acid level took 1.24 min (Mash took 2.59 min) with similarly high top 5 and top 10 recall (99.7% and 98.5%, respectively) against brute-force Mash and Blastp-based AAI. Note that the difference in run time will be much larger between Mash and GSearch as the number of fungal database genomes increases in the future, as also exemplified above for the bacterial genomes.

### Combining the three-step framework for prokaryotic genome classification and comparison with standard tool GTDB-Tk

To evaluate the accuracy of classification/searching results based on the complete 3-step framework of GSearch, we compared the best neighbors found by GSearch with brute-force ANI (estimated by FastANI) and GTDB-Tk, a standard for microbial taxonomy classification these days that is phylogenetic-tree– and alignment-based. As mentioned above, when the query genome does not find a match in the database better than ANI > 78%, corresponding to ProbMinHash distance 0.9850, the nucleotide-level graph is abandoned, and the amino-acid level is used instead; similarly for no match better than 52% AAI, GSearch switches to the universal gene graph (Figure 2g). The overall running time to classify 1000 prokaryotic genomes of varied levels of taxonomic novelty compared to the database genomes on different computing platforms is showed in Table S5. On a 24-thread Linux node with Intel Xeon Gold 6226 CPU, it took GSearch a total of 5.85 minutes while it took 19.49 minutes on an intel Core i7 laptop (2017 release) CPU personal laptop (6.02 minutes on the most recent ARM64 CPU laptop). Classifying 1000 genomes using GTDB-Tk took 5.91 h on the same Linux node with 24 threads (memory requirement was ∼328GB) while MASH took 53.7 min for 1000 genomes using 24 threads. We were not able to assess Dashing or BinDash for this analysis because they do not provide proteome (amino acid) level implementation.

In terms of classification accuracy, all query genomes that had a best match higher than 78% ANI against the GTDB database genomes (i.e., a match at the same or closely related species, 699 out of the total 1000 queries) were identically classified by GSearch, GTDB-Tk and FastANI (Supplementary file 1, only 100/699 are shown for simplicity), which means species-level classifications are highly consistent. For the remaining 301 genomes that did not have same or closely related species-level matches, for 266 of them (or 87.1%), GSearch also provided the same classification with GTDB-Tk but several inconsistencies were observed for 39/301 genomes (Figure 3e). Specifically, we noticed that for GTDB-Tk, which relies on RED values and tree topology, several genomes (n=14) were still classified at the genus level even though the AAI value against the best database genome in these case was below 60% (typically, genomes assigned to the same genus show >65% AAI(46)), and some genomes (n=16) were still classified at the family level but not at the genus level even though their best AAI value was above 65%. Similarly, several genomes (n=9) were classified at the order level but not family level even though their best AAI value was above 52%. Therefore, high consistency was overall observed between GSearch and GTDB-Tk assignments, and the few differences noted were probably associated with contaminated (low quality) MAGs or taxonomic inconsistencies, which was challenging to assess further, and/or the peculiarities of each method. Since ProbMinHash distance correlated well with blastp-based AAI after transformation in the range of AAI values between 52% and 95%, the classification results were always consistent with AAI-based classification using previously proposed thresholds. For example, best matches at AAI 65% ≥ AAI were classified in the same genus by GSearch and blast-based AAI, and best matches of 52% < AAI < 65% were typically classified in the same family (but different genus within the family).

### Database split for large genome databases

For large databases (for example, >1 million bacterial genomes), the graph building and requesting step could require a large amount of memory (due to the larger k-mer space) that is typically not available in a single computer node. We therefore provide a database split solution for such large databases. The average database building time on each node for each piece of the database after the splitting step scales linearly with increasing nodes/processors (Figure S7a) and requires much less memory (1/n total memory compared to when building in one node, where n is the number database pieces after splitting; for GTDB v207 nucleotide graph building and n=5, it will be only 28.3/5=5.66 GB). The searching time scales sub-linearly with increasing number of nodes (Figure S7b) but offers the advantage of a reduced memory footprint with respect to the single-node search. The top 10 best neighbor by splitting the database were exactly the same as the non-splitting strategy (Supplementary file 2). Note that without multi-node support (e.g., run database build sequentially), database build time is nearly the same with non-split strategy, but memory requirement is only 1/n (GTDB v207, 28.3GB/5=5.66GB at nucleotide level and 27.7GB/5=5.54GB at amino acid level), even though total request time will be longer (time*n in Figure S7(b)). However, since the request step is very fast, requiring only 1/n memory compared to non-split strategy, overall runtime is still short with the database split approach even for a relatively large number of genomes. The database split strategy is especially useful when memory requirement is not satisfied on the host machine for larger genome database (e.g., millions of genomes).

## Discussion

A popular way to assess genetic relatedness among genomes is ANI, which corresponds well to both 16S/18S rRNA gene identity and DNA-DNA hybridization values, the golden standards of fungal and prokaryotic taxonomies (7). However, the number of available microbial genomes has recently grown at an unprecedented speed. For example, there are 30% more (new) species in NCBI/RefSeq 2022 compared to 2023, and the number of bacterial species represented by genomes alone is expected to surpass 1 million soon. Therefore, the traditional way that blast-based ANI or faster k-mer-based implementations (e.g., Skani, FastANI, Mash or Dashing 1/2) are applied as an all vs. all search strategy (brute-force) does not scale because the running time grows linearly with increasing number of query genomes and/or genomes in the database. Phylogenetic approaches based on quick (approximate) maximum likelihood algorithms and a handful of universal genes as implemented –for example-in GTDB-Tk could be faster than brute-force approaches but are often not precise and require a large amount of memory for the querying step (11) while the database building step could take several weeks of run time because the underlying multiple sequence alignment of the database genomes is computationally intensive. Further, approaches that reply on k-medoid clustering to avoid all vs. all comparisons could be sometimes trapped into local minima because of arbitrary partitioning of database genomes into clusters, a known limitation of these methods (13). GSearch effectively circumvents these limitations by combining new k-mer hashing-based probabilistic data structures for fast computation of genomic relatedness among genomes (i.e., ProbMinHash, SuperMinHash and/or SetSketch) with a graph based nearest neighbor search algorithm (HNSW). Accordingly, GSearch is at least an order of magnitude faster than alternative approaches (e.g., Mash, Dashing, Sourmash, GTDB-Tk) for the same purposes and will be even faster as the number of database genomes grows. Therefore, GSearch solves an important challenge associated with the tasks to search and/or classify microbial genomes and will serve well these tasks for years to come based on the current speed in the increase of the number of genomes that become available.

To the best of our knowledge, no current tools have combined two sub-linear algorithms and thus efficiently search very large genome databases while maintaining high accuracy in calculating ANI/AAI among the genomes, and hence identifying the true best neighbors. Several sub-linear algorithms such as Sequence Bloom Tree or COBS (cross-over between an inverted index and Bloom filters) can do sequence to sequence (or genome to genome) search but do not provide ANI/AAI values, which are key for microbial genome search and classification (Table 1). Accordingly, these tools are generally less accurate than the ANI/AAI approach. GSearch can handle several million of microbial genomes on a small-to-average computer cluster, or even personal laptop (depending on the database size), since the created (dumped) database file size is proportional to the total number of genomes in database for fixed sketch size and graph parameters, and generally rather small. Specifically, with one million genomes, the dumped file size will be 5.9GB*20=118 GB (currently, there are ∼60K in the GTDB database, creating a database file size of 5.9GB). With the SetSketch option, database file size will only be 0.5GB for the GTDB database, 2.5G for the entire RefSeq genome databases (∼318K genomes, 2 Terabytes), and 9.8GB for one million genomes. Further, due to the nature of the graph based NNS algorithms, there is no need to build the entire database at once, but the database can be split it into smaller pieces, as exemplified above and depending on the computational resources available. For a modern laptop with 16 GB memory, a database on one million species can be split into 10 pieces, so the dumped file for each piece will be only 11.8 GB, which can be loaded into memory, and then collect the results from each piece within an approximate total running time of 30 minutes (for querying 1000 genomes against 0.1 million database genomes for each of the ten parts, which should take about 3 minutes). With this logic, a computing node with 24 threads and 256 GB of memory available can easily deal with 20 million bacterial database genomes and billions of bacterial genomes if using SetSketch. This represents a substantial improvement compared to existing tools for the same purposes. This database split/partition idea of HNSW, or other graph-based NNS libraries, has been successfully applied in several industrial-level applications for other purposes (31,32). We also want to point out that the size of the compressed sketches (database file) from SetSketch is comparable to other space efficient algorithms such as HyperLogLog (with b=2, Setsketch asymptotically corresponds to HyperLogLog), ExtendedHyperLogLog (47), HyperLogLogLog (48) and UltraLogLog (49) (25% more space efficient than HyperLogLog). The Shannon entropy of SetSketch we implemented is 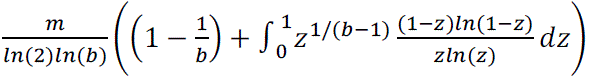, where m is the number of registers, b is the key parameter of SetSketch. For b=0.0005 that we used, the entropy is 13.24 per register (divided by m). Therefore, compared to an uncompressed register size of 16 bits, there is still some room for improvement for SetSketch. Theoretically, a 17.25% reduction in space can be achieved but would add additional implementation complexity. For any HyperLogLog-like algorithms (e.g., SetSketch, UltraLogLog), a theoretical limit for sketch size is *O*(ϵ^8’^ + *log*(*n*)), or slightly worse but easy to implement (50,51), where n is the number of elements/k-mers while ɛ is error. This means that we cannot improve further without losing accuracy as database size grows unless we rely on database split strategy. SetSketch shares this property with HyperLogLog and thus we cannot improve space further without theoretical breakthroughs.

Another distinguishing aspect of GSearch (tohnsw module) is the speed and flexibility in building reference databases. Users could build reference databases (graphs) for any number and type (e.g., prokaryotic vs. viral) of genomes. The high efficiency in building graphs allows users to also test and optimize the key parameters of the graph, the M (maximum allowed neighbors) and ef_construct (width of search for neighbor during building) parameters. For any given database size, M and ef_construct determine the quality of the graph and graph build speed. Small M and ef_construct may lead to frequent traps in local minima and thus, low recall while large M and ef_construct may lead to slow speed without proportional improvement in recall (Table S7). Therefore, there is a tradeoff between accuracy and speed that should be evaluated first. However, for most users this task would not be necessary because they will work with pre-built databases such as those provided here. Also, GSearch provides an option to add new genomes to the database without the need to rebuild the database from scratch, making it convenient and very fast to expand the current genome database as needed.

GSearch could also be applied to whole metagenome search and identification of the most similar metagenomes in a series because the relatedness among metagenomes can be estimated in a similar way to genomes using ProbMinHash as implemented –for example-in the HULK software, which relies on the ProbMinHash algorithm in a streaming fashion (52). However, to allow for sketching of much larger sequence files such as whole metagenomes simultaneously to build the HNSW graph, weighted k-mer (e.g., ProbMinHash) approaches require much larger memory than unweighted approaches due to the larger number of total k-mers (including duplicates). Probabilistic or approximate counting of k-mer abundances such as Count-Min sketch can be used instead for k-mer counting for metagenome search and/or when the computer memory is limited (53,54), or a combination of Count-Min Sketch with HyperLogLog sketch (55). Similarly, we could seamlessly replace ProbMinHash with another relatedness algorithms should such algorithms become available and has advantages in terms of speed and/or precision, or more importantly, has the locality sensitive hashing property for nearest neighbor search. For example, the SuperMinHash option is now provided for its high accuracy over traditional MinHash but not faster (37), or other newer implementations such as the One Permutation MinHash with Optimal Densification (OPMHOP) for estimating the unweighted Jaccard index, due to its speed (only one hash function, average case O(n+m)) and locality sensitive hashing property (via optimal densification or faster densification)(35,36) (Table S1 and S2). Note that the Bottom-K sketch implementation in Mash or FracMinHash in Sourmash use just one single hash function or a much smaller number of hash functions required by that of classic MinHash and thus lose the locality sensitive hashing property and thus we cannot use it for nearest neighbor search purposes (36). Further, BagMinHash (56) or DartMinHash (57) for weighted (but not normalized) Jaccard index can also be used in place of ProbMinHash (also LSH). Since the number of genomic distance computations is O(log(N)) in GSearch, the computational time for estimating genomic distance for a pair of genomes is not a major factor in overall computational speed because log(N) is always a small number. Accuracy in the genomic distance estimate is relatively more important in GSearch and the main reason that ProbMinHash is the default option. Related to this, ANI as currently implemented –for instance-in FastANI is not appropriate for this function because it is not metric; that is, for the FastANI distances calculated among three genomes A, B, and C, (A,B) + (B, C) is not necessary larger than (A,C) and ANI(A,B) is not equal to ANI (B, A)^15^. Similarly, mutation rates (or ANI) estimated by FracMinHash, CMash and related tools (e.g. Sourmash or skani calculated ANI) are also not metric (58–60). To solve this “metric” problem, a norm adjusted proximity graph (NAPG) was proposed based on inner product and it shows improvements in terms of both speed and recall using non-metric distances (61). This could be another direction for further improving the speed and recall of GSearch. The three options we provided as part of GSearch to estimate Jaccard-like index are all metric, which ensures neighbor diversity when building the graph, and they are equally applicable to any microbial genome, including viral and fungal genomes.

K-mer-based methods for genetic relatedness estimation such as ProbMinHash have lower accuracy between moderately-to-distantly related genomes compare to alignment-based tools (see Supplementary Note 2 and 4 for further discussion). Our empirical evaluation showed that this relatedness level, for nucleotide searches, is around 78% ANI and 52% AAI for the amino-acid searches (e.g., ProbMinHash distances do not correlate well with blast-based ANI and AAI at and below these levels). To circumvent this limitation, we designed a 3-step framework as part of GSearch to classify bacterial genomes that show different levels of novelty compared to the database genomes, with high accuracy. This framework included a search at the universal gene level for deep-branching genomes that are novel at the phylum level (AAI < 52%), for which searching at the entire proteome level is less accurate. Recently, methods that employ k-mers that allow mismatches, that is, spaced k-mers have shown promise in accurately estimating genomic relatedness even among distantly related genomes with gains in speed (62). To apply spaced k-mers to entire genomes, the recently developed “tensor sketch” approaches could be explored in the future to simplify the distance computation for distantly related bacterial and viral genomes (63) instead of relying on the mentioned 3-step framework. In the meanwhile, the ProbMinHash approach, essentially a Jaccard-like distance estimation via weighted and normalized MinHash for k-mers, is highly efficiently, and, importantly, can effectively deal with incomplete genomes or genomes of (drastically) different length, a known limitation of traditional Mash-based methods (22). Comparing genomes of different length is not uncommon, e.g., bacterial genome sizes can differ by more than ten-fold, as can be the case between MAGs of different level of completeness or when searching a short sequence (e.g., a bacteriophage genome) against a large genome collection (e.g., the whole viral genome database). Our own analysis showed that ProbMinHash is robust down to 50% completeness level (Table S12), which is also the most commonly used standard for selecting MAGs of sufficient/high quality for further analysis and reporting (64). ProbMinHash is also robust for genomes with repeats or gene duplications (e.g., fungal genomes) due to the k-mer weighting step by weighted MinHash, a property not shared by simple MinHash.

In general, the genome relatedness estimated, or best database matching genomes identified, by GSearch were highly consistent with blast-based ANI/AAI results or phylogenetic placement of the genomes using GTDB-Tk, particularly for query genomes with close relatives in the database related at the species or genus levels (Supplementary file 1, Figure 3e). For more distantly related query genomes relative to database genomes, classification results of GSearch showed some differences with GTDB-Tk. These differences were not always possible to assess further for the most correct genome placement but could be due, at least partly, to the incompleteness and/or contamination of query or/and database genomes, which renders the resulting concatenated alignment of universal genes used by GTDB-Tk unreliable (64,65), as only a few amino-acid positions per gene are used in the final alignment. In contrast, the AAI and ProbMinHash approaches should be more robust to changes of a small number of genes because the entire proteome is considered.

Graph-based NNS methods achieve good performance compared to tree based and locality-sensitive hashing (LSH) methods or space partitioning methods (33). Building a HNSW graph relies on proximity of the database elements; thus, if the distances among database elements, in our case genomes, cannot be effectively estimated, the navigation of graph becomes less efficient (e.g., gets trapped/lost in local minima). This is especially problematic for highly sparse/distantly related and diverse datasets, like the viral genome database, in which two phage genomes could often share very little genomic information (k-mers), and thus the edges to choose from during graph navigation are not accurate estimations of the relatedness of the corresponding genomes. This is confirmed by our own results when using nucleotide-level search to build the viral genome graph. Hence, the amino acid level will be much more robust for viral genomes and is the recommended level to use. Finally, recent advancements in proximity graph building could further reduce database building time from O(N*log(N)) to O(N*c) (where c is a constant independent of N, number of genomes) and/or achieve better search performance than existing approximate proximity graph (APG) alone by combining locality sensitive hashing (LSH) with approximate proximity graph (66) (or APG, e.g. HNSW is a APG) to have LSH-APG. This approach essentially reduces the number of points/genomes to be compared during graph building and will be explored in future versions of GSearch to provide additional speed-up and/or robustness for graph database building. The HNSW graph, and graph-based K-NNS in general, can also be further improved by adding shortcut edges and maintaining a dynamic list of candidates compared to a fixed list of candidates used by default (67). Graph reordering, a cache optimization that works by placing neighboring nodes in consecutive (or near-consecutive) memory locations, can also be applied to improve the speed of HNSW (68). We also want to mention that in our HNSW Rust implementation, a memory map was also implemented, which will solve the memory limit problem when building large graph with billions of data points or genome. Another direction to further improve GSearch could be the use of Graphics Processing Unit (GPU) instead of CPU because GPUs are more efficient in handling matrix computations and machine learning tasks (69). We will explore these options in future versions of GSearch.

Finally, GSearch provides a general framework/idea to create a new data structure: combining probabilistic data structures (e.g., MinHash, HyperLogLog) with graph-based nearest neighbor search algorithm that should be applicable not only to genomes but also text or document searching purposes more broadly. For example, the MinHash algorithm can be applied to document (e.g., website, text) similarity estimation, and be combined with HNSW. Further, we can hash strings to approximate edit distance (a metric distance) to avoid expensive dynamic programming (70) via order MinHash (71). We reimplemented order MinHash in our ProbMinHash library, and it can be easily applied to do fast DNA string (not fragmented genomes) or general string search.

To summarize, GSearch, based on MinHash-like algorithms and HNSW, solves a major current challenge in search and classification of microbial genomes due to its efficiency and scalability. Both the Densified MinHash and HNSW are approximating the theoretical optimal limits of similar algorithms in terms of speed and accuracy. GSearch will serve the entire microbial sciences for years to come since it can be equally applied to fungal, bacterial, and viral genomes, and thus offer a common framework to identify, classify and study all microbial genomes, at the million-to-billion database genome level.

## Data and Code availability

All the mentioned pre-built database for bacteria, fungi and phage genomes can be found at: http://enve-omics.ce.gatech.edu/data/gsearch. Code can be found here: https://github.com/jean-pierreBoth/gsearch or here: https://gitlab.com/Jianshu_Zhao/gsearch. Scripts for reproducing the major results can be found here: https://github.com/jianshu93/gsearch_analysis.

## Author Contribution

J.Z, and K.K designed the work, J.Z and J.P-B wrote the code (Genome part and algorithm part respectively), J.P-B implemented the Rust libraries of Kmerutils, Probminhash and Hnswlib-rs. L.M.R provided constructive suggestions. J.Z did the analysis and benchmark. J.Z and K.K wrote the manuscript.

## Acknowledgment

This work was supported, in part, by the US National Science Foundation (Award No 1759831 and 2129823 to KTK). We want to thank PACE (Partnership for Advanced Computing Environment) at Georgia Tech for providing computing resources. We want to thank Kenji Gerhardt for helpful discussions on benchmarking against traditional ANI/AAI based tools and Chirag Jain for discussions on MinHash and bottom-k sketch implementation. We also want to thank Otmar Ertl for discussion on SetSketch Shannon entropy and implementation of SuperMinHash/ProbMinHash, Xiaofei Zhao for discussions on mergeability of One Permutation MinHash with Optimal densification, and Dingyu Wang for discussion on τ-GRA relative variance with respect to HyperLogLog. Finally, we would like to thank two anonymous reviewers for their suggestions, which help improve the manuscript substantially.

