## Supplementary_materials for "GSearch: Ultra-Fast and Scalable Microbial Genome Search by Combining K-mer Hashing with Hierarchical Navigable Small World Graphs"

**This file contains the following:**

**1. Supplementary Figures, page 2-11**

**2. Supplementary Tables, page 12-20**

**3. Supplementary Methods & Materials, page 21-27**

**4. Supplementary Notes, Page 28-35**

### Supplementary Figures

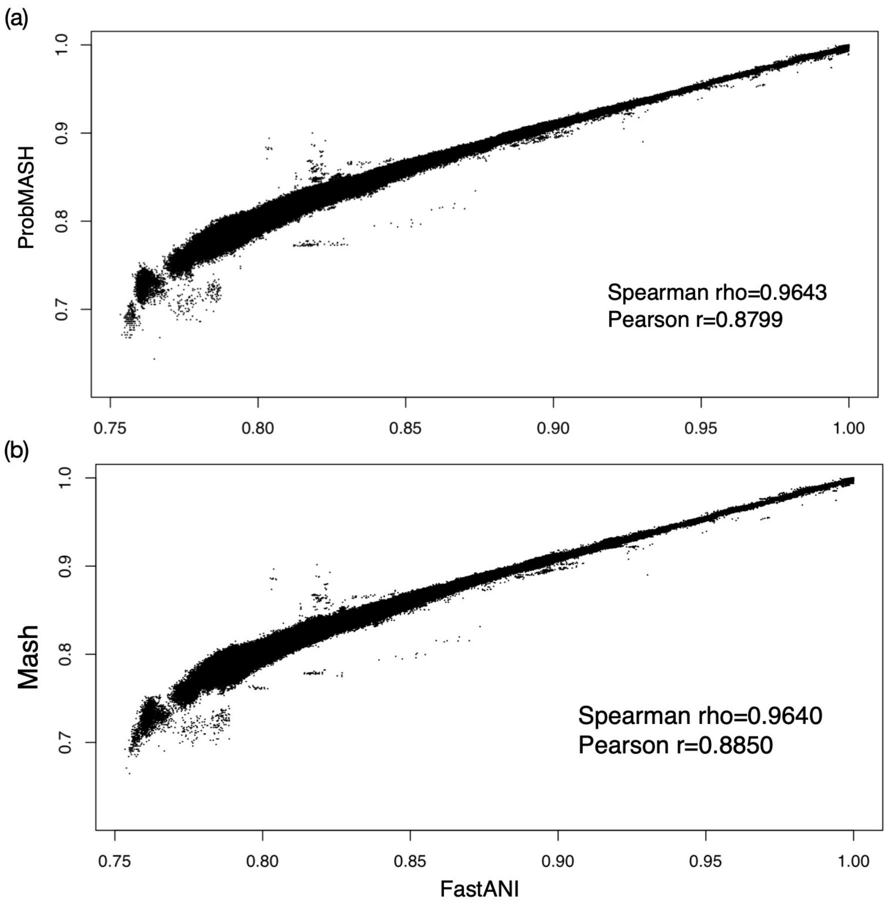


**Figure S1**. Relationship between FastANI and ProbMASH (a) and FastANI and MASH (b) distances for closely related genomes. Both Spearman and Pearson correlation significance tests showed $p<0.001$ in (a) and (b). ProbMASH is the transformed ProbMinHash distance according to the MASH distance formula (See Materials and Methods, ProbMinHash section).

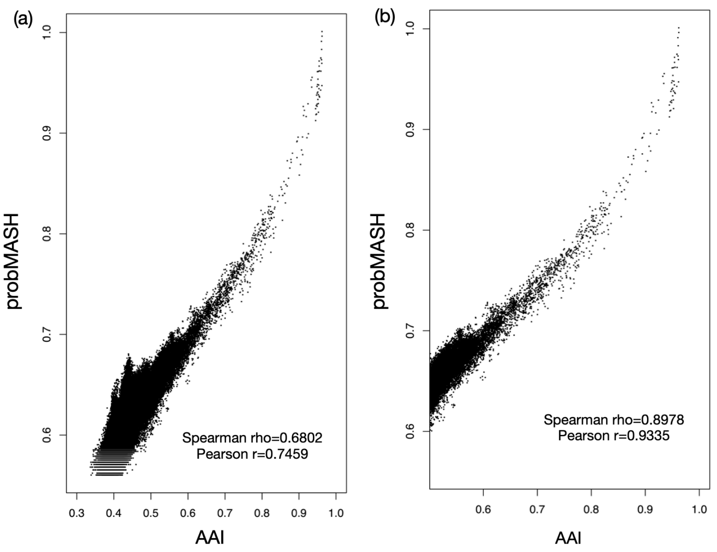


**Figure S2**. Correlation between AAI (entire proteome) and ProbMASH distance for (a) all AAI values and (b) AAI values between 0.52 and 0.95 among 2000 genomes randomly selected from the GTDB v207 database. Both Spearman and Pearson correlation significance test showed p<0.01.

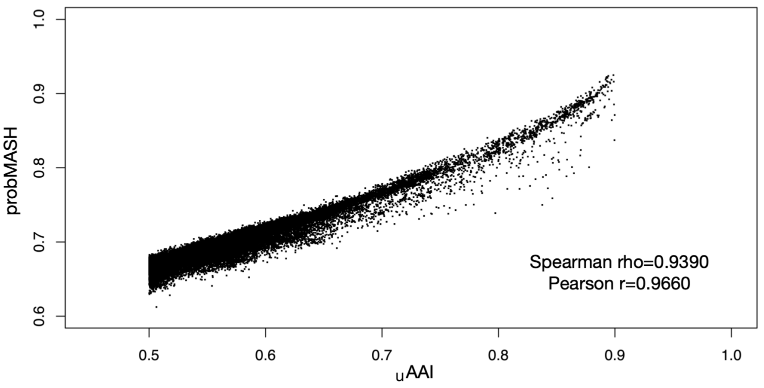


**Figure S3**. Correlation between AAI (universal gene set, or uAAI) and probMASH distances for AAI values between 0.5 and 0.9 among 2000 genomes randomly selected from the GTDB v207 database. Spearman and Pearson correlation significance test showed $p<0.01$.

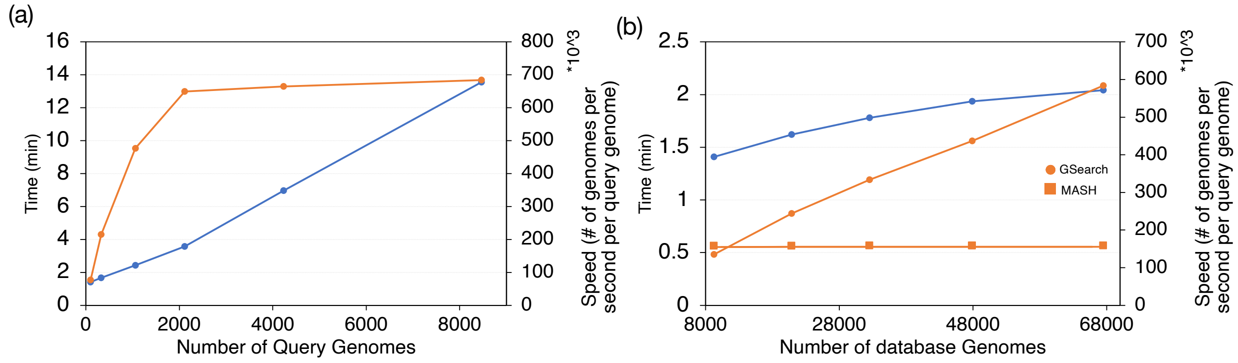


**Figure S4**. Performance of GSearch search step. (a) Search time (blue) and speed (orange) for an increasing number of query genomes against a fixed database of 67503 genomes and, (b) search time (blue) and speed (orange) for an increasing number of database genomes and a fixed number of 1059 query genomes. Search speed is defined as the number of database genomes searched, on average, per query genome per second. The orange line with squared data points represents Mash, whose speed was constant for any database size for 1059 query genomes, for comparison.

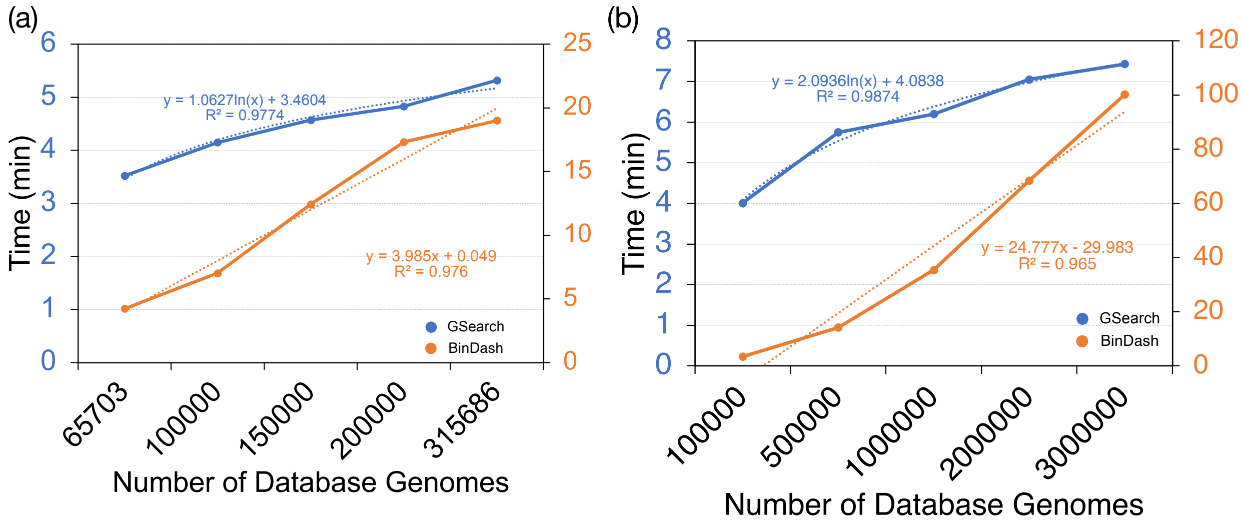


**Figure S5**. GSearch request speed comparison with BinDash software for (a) all bacterial genomes (~318K, 2T) and (b) all viral genomes (~3 million) as database. 8,466 and 10,000 query genomes were used, respectively, for this analysis. Note that GSearch and BinDas are plotted in different axes, left y and right y axis, respectively. 24 threads were used. Sketch size m=10^5 was used to achieve similar accuracy with ProbMinHash.

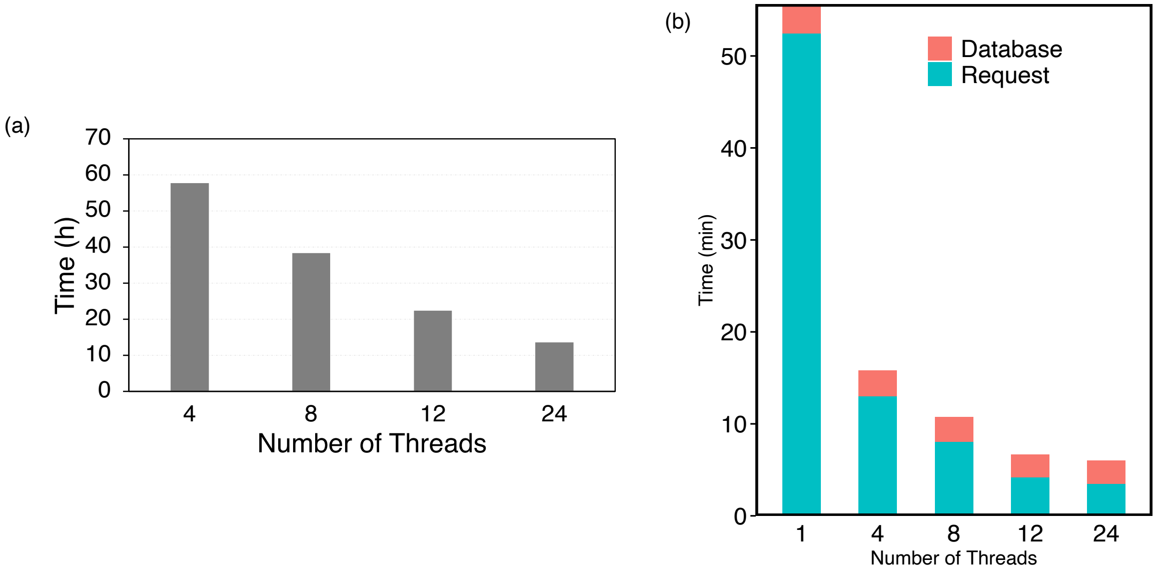


**Figure S6**. Database build time for the IMGVR phage species database (935,122) at the amino acid level (a), and total search (request) time for 10,000 query phages (b) against this database. Database size was about 15.8 GB, and thus loading database takes a substantial time of the total searching time.

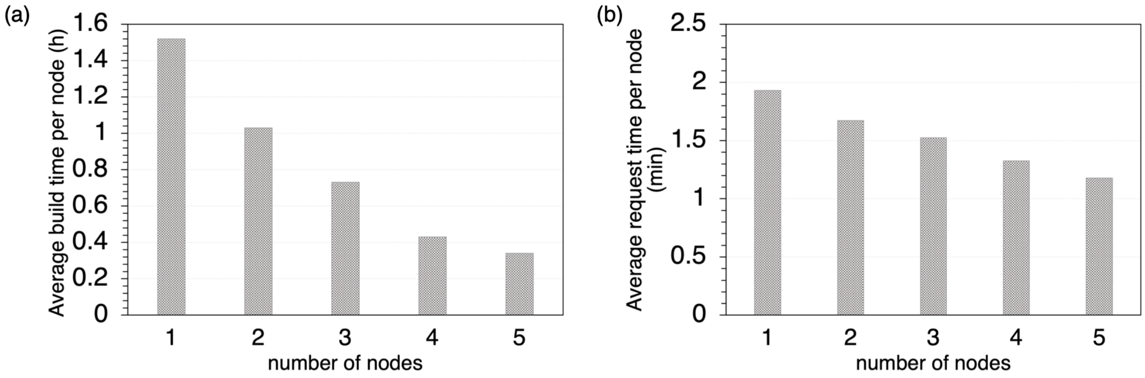


**Figure S7**. Database build time vs number of nodes or number of database pieces (maximum 5) for the split strategy (a), and request/search time of 100 bacterial genomes (nt) against the split databases (b). Each node was responsible for running a piece of the full database. Average time was calculated by averaging total time across all nodes. The entire NCBI/RefSeq database (318K genomes) were used for testing the split strategy. For each node, 24 threads were used for building and searching steps.

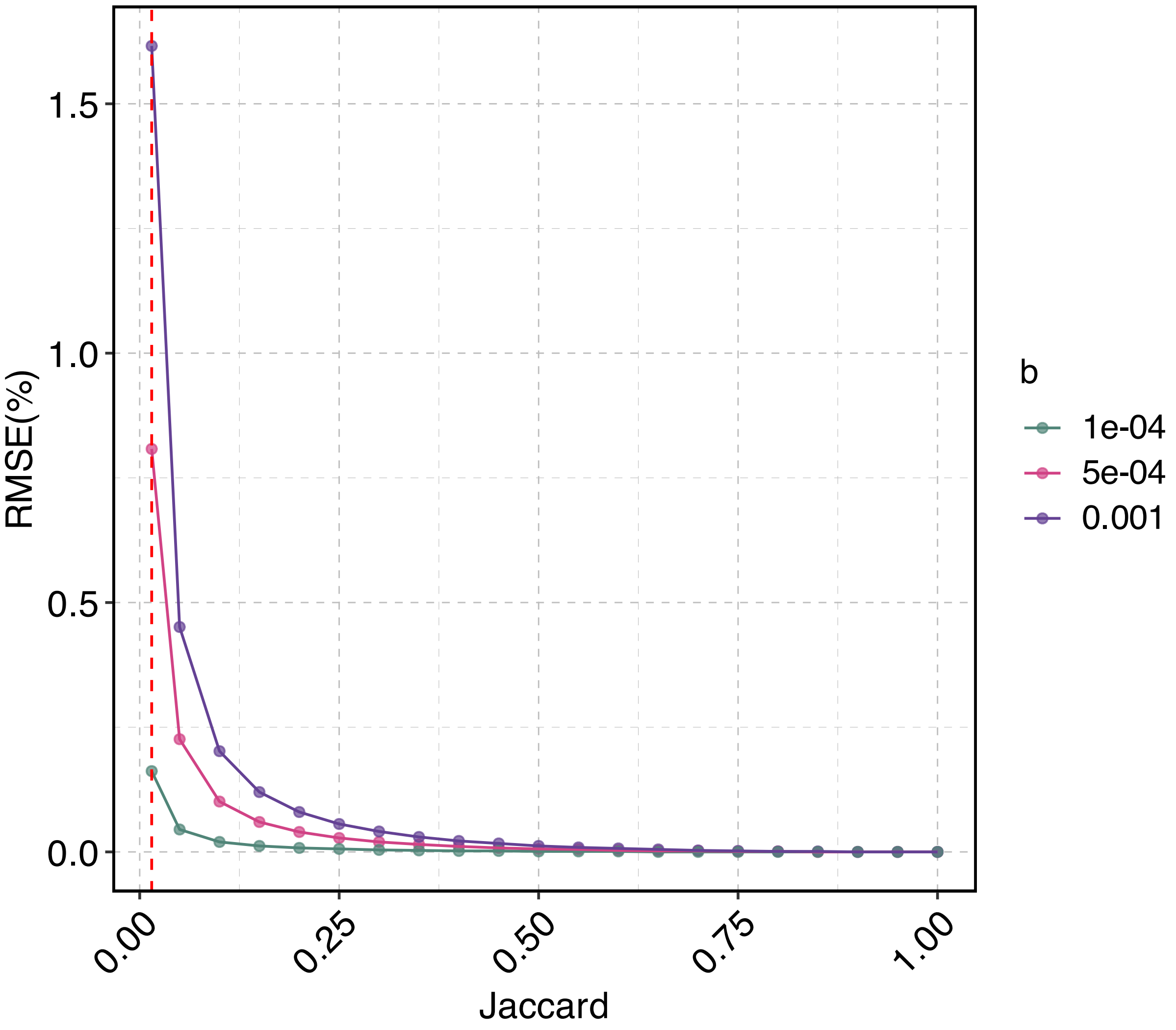


**Figure S8.** Relative Mean Square Error (RMSE) (MSE divided by true Jaccard) of SetSektch LSH algorithm for estimation of Jaccard index with respect to true Jaccard index for different values of parameter b. Note that changing b requires changing m in SetSektch. Three b values were used, 0.001, 0.0005, 0.0001 corresponding to m=4096,6144 and 8192 respectively. The red dashed line indicates J=0.015 with RMSE 0.81%, corresponding to ANI 77.99%, with variation of ANI from 77.94% to 78.04% for the case that b=0.0005 and m=6144.

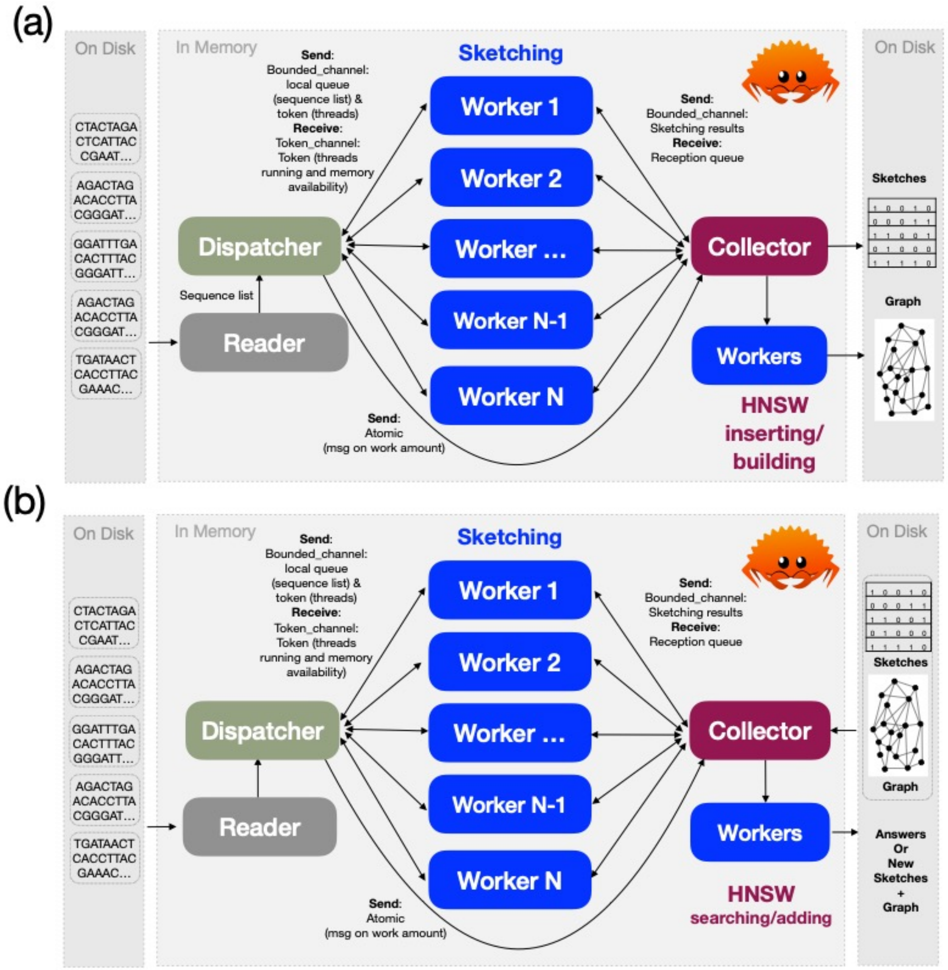


**Figure S9**. Parallel and Concurrent computing model of GSearch tohnsw (a) and request or add (b) modules. Standard dispatcher and worker model was used with Reader and Collector to handle file input, and collecting results from sketching workers and initializing HNSW graph building/searching workers. Tohnsw/add module will dump sketching results and graph to disk while searching module will load prebuilt sketching and graph files and then dump answers for the query genomes to disk. Note that the number of Workers for HNSW building/searching/adding is to use all available computing cores while the number of Workers for sketching can be controlled to balance memory consumption and sketch speed. The concurrent model was based on Rayon library and communication among threads was based on Crossbeam library.

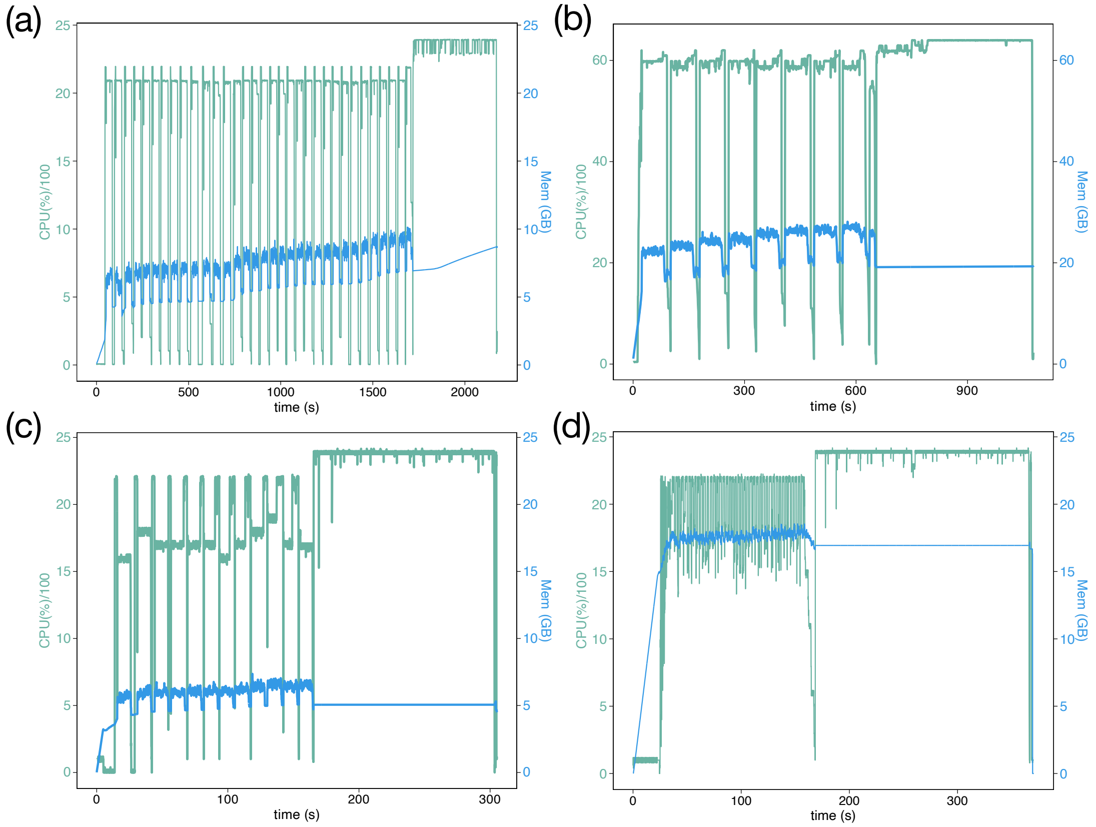


**Figure S10.** Parallel efficiency and memory consumption of GSearch tohnsw and request modules (ProbMinHash) at the nucleotide level. (a) tohnsw build module real-time CPU usage (left axis) and memory consumption (right axis) for the GTDB v207 (65,703 genomes) database using 24-thread node. (b) The same with (a) but on a 64-thread node. (c) Real-time CPU usage and memory consumption for searching 8,466 genomes against the GTDB prebuilt database in (a) on a 24-thread node. (d) Real-time CPU usage and memory consumption for searching 8,466 genomes against entire NCBI/Ref_Seq genomes (~318K). Two phases in all figures are sketching and graph building/searching, respectively.

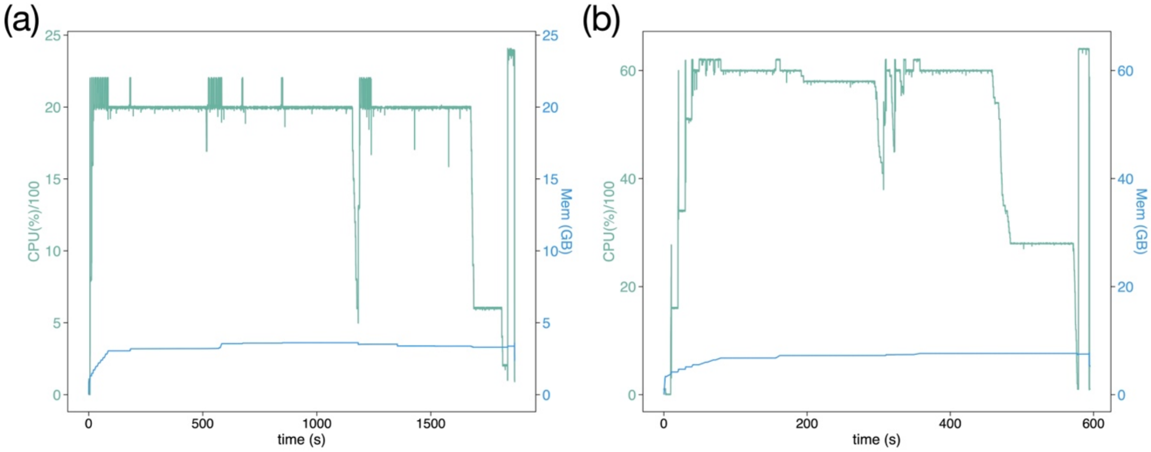


**Figure S11**. Parallel efficiency and memory consumption of GSearch request modules (SetSketch) at nucleotide level. (a) 8,466 query genomes against the GTDB v207 database (65,703 genomes) using 24 threads. (b) 8,466 query genomes against NCBI's RefSeq database (~318K genomes) using 64 threads.

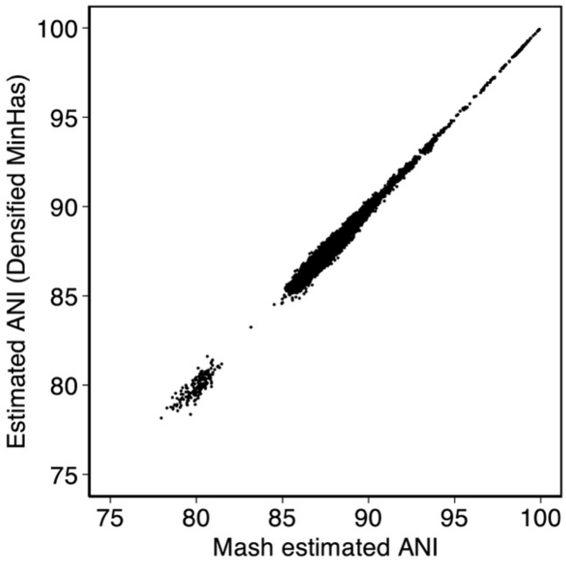


**Figure S12**. Correlation between ANI estimation from our implementation of densified MinHash (optimal densification) and Mash bottom-m implementation. We use sketch size m=10^5 to improve accuracy while 1.2*10^4 was used in Mash. Our implementation, as also reported in BinDash, is at least 10 times faster than Mash.

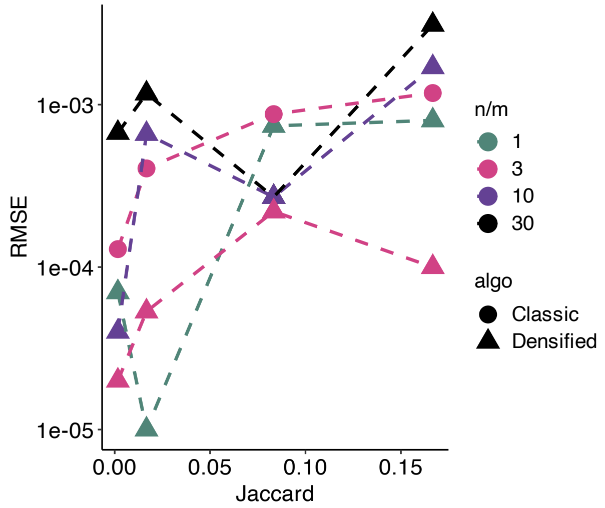


**Figure S13**. Rooted Mean Square Error (RMSE) of Densified MinHash and Classic MinHash theoretical RMSE ($RMSE = \frac{\sqrt{J*(1-J)}}{\sqrt{m}}$) for small Jaccard index (more challenging than large Jaccard index), where m is the sketch size (or number of registers). We use set size (number of elements in the set) n=300,000 for all cases. Jaccard index is 0.001667, 0.01667, 0.0833, 0.1667 respectively for each point in x axis.

### Supplementary Tables

**Table S1.** Comparisons of MinHash-like and HyperLogLog algorithms. Bold face results were obtained as part of this study (via gsearch --algo prob/super/hll option); remaining results are from the literature. $J_p$ is closer to real Jaccard index than $J_w$ despite both algorithms consider k-mer weights.

| | Weighted element | Set Size bias | Speed (More is better) | Space (Less is better) | Mergeability[10] | Approximated Jaccard-like index |
|---|---|---|---|---|---|---|
| Classic MinHash[1] | ✗ | ✓ | ★★★★ | ☆☆☆☆ | ✓ | J |
| **Densifed MinHash (with OD[2,11])** | ✗ | ✓ | ★★★★★★ | ☆☆☆ | ✗ | J |
| **SuperMinHash[3]** | **✗** | **✓** | **★★★** | **☆☆☆** | **✓** | **J** |
| **ProbMinHash (default)[4]** | **✓** | **✗** | **★★★★★** | **☆☆☆** | **✓** | **$J_p$** |
| BagMinhash[5] | ✓ | ✓ | ★★ | ☆☆☆☆☆ | ✓ | $J_w$ |
| DartMinHash[6] | ✓ | ✓ | ★★★★ | ☆☆☆☆☆ | ✓ | $J_w$ |
| HyperLogLog[7] | ✗ | ✗ | ★★★ | ☆☆ | ✓ | J |
| HyperLogLogLog[8] | ✗ | ✗ | ★★ | ☆ | ✓ | J |
| **SetSketch[9]** | ✗ (✓)[12] | **✗** | **★★★** | **☆☆** | **✓** | **J** |
| UltraLogLog | ✗ | ✗ | ★★ | ☆ | ✓ | J |

[1]Broder (1997), [2]Shrivastava (2017), [3]Ertl (2017), [4]Ertl (2020), [5]Ertl (2018), [6]Christiani (2020), [7]Flajolet et al. (2007), [8]Karppa and Pagh (2022), [9]Ertl (2021)
[10] which means that adding an element or taking the union of multiple subsets can be performed in sketch space, important feature for large scale application when new data need to be added to existing sketches or distributed environment.
[11]OD indicates optimal densification. Since it is not mergeable, it is difficult to be applied in distributed environment.
[12]SetSketch can be applied in a weighted fashion but not in this paper

**Table S2.** Theoretical comparison of estimator variance (relative mean standard error) of MinHash-like algorithms for Jaccard index, where m is the number of registers (sketch size). We use the same m for all tools evaluated expect for SetSketch, which requires a smaller m to have similar accuracy. J or $J_p$ represents Jaccard index or $J_p$ respectively.

| | RMSE |
|---|---|
| Classic MinHash | Sqrt(J*(1-J)/m) |
| ProbMinHash | sqrt($J_p$*(1-$J_p$)/m) |
| SuperMinHash[1] | $sqrt(\frac{J(1-J)}{m}(1-\frac{\sum_{l=1}^{m-1} l^u((l+1)^u+(l-1)^u)-2l^u}{(m-1)^u m^u (u-1)}))$ |
| SetSketch[2] | [sqrt(1/m), sqrt(1.08/m)] |
| Dashing (original&Improved)[3] | sqrt(1.079/m) + union/intersection error > sqrt(1.079/m) |
| Dashing (MLE)[4] | sqrt(1.074/m) |
| Densifed MinHash[5] | sqrt(1/m) |
| $\tau$-GRA ($\tau$=0.8898)[6] | sqrt(1.075/m) |

[1]The second long term in the variance is always smaller than 1, thus SuperMinHash RMSE is always smaller than classic MinHash, the smallest MinHash RMSE known until today; [2]As b is close to 1, RMSE is close to lower bound; [3]Both the original and improved estimator need correction at small cardinality. Dashing2 has the same variance with SetSketch because it reimplemented SetSketch; [4]MLE is Maximum Likelihood Estimator, theoretically optimal for HyperLogLog sketches but much slower; [5]One Permutation MinHash with Optimal Densification, variance goes to 0 much faster than classic MinHash/SuperMinHash/ProbMinHash as m increases, see Shrivastava (2017). Note that the variance is larger than classic MinHash or ProbMinHash for any J because J(1-J) is always smaller than 0.25; [6]Generalized Remaining Area with $\tau$=0.8898 has the lowest variance. Detailed analysis of RSME for Tau-GRA can be found in Pettie and Wang (2022)

**Table S3.** Comparisons of database size, running time for ProbMinHash, SuperMinHash and SetSketch with Dashing and BinDash. Results are based on searching 8,466 query genomes against all NCBI/RefSeq genomes (~318K) on a 24-thread machine. Times reported are average values from 3 runs.

| | Database size | database loading time | Sketching & searching | Total Wall time |
|---|---|---|---|---|
| GSearch(ProbMinHash) | 29G (sketch + graph) | 57.2s | 8 min 23 s | 9 min 20 s |
| GSearch(Densified MinHash) | 9.7G | 27.2 s | 4 min 12s | 4 min 40 s |
| GSearch(SuperMinHash) | 25G (sketch + graph) | 41s | 14 min 13 s | 14 min 54 s |
| GSearch(SetSketch) | 2.5G (sketch + graph) | 9.1s | 12 min 14 s | 12 min 23 s |
| Dashing 1 | 2.7G (sketch) | ~61s | 27.5 min | 28.5 min |
| Dashing 2 (SetSketch) | 4.7G (sketch) | ~61s | 27.5 min | 28.5 min |
| Skani[1] | 30.5G | - | 64.42 min | 64.42 min |
| BinDash | 6.2G (Sketch) | 16s | 21.4 min | 21.4 min |

[1] for ~8000 query genomes, skani memory requirement is larger than 60G the largest among all

**Table S4.** Comparisons of database size, running time for GSearch ProbMinHash, SuperMinHash and SetSketch for IMG VR virus database. Results are based on searching 10,000 qery genomes against all IMG_VR v4 genomes (~3 million) on a 24-thread machine. With 3 million genomes, graph file loading dominates the running time for GSearch. Times reported are average values from 3 runs.

| | Database size/memory | database loading time | Sketching & searching | Total Wall time |
|---|---|---|---|---|
| GSearch(Prob MinHash) | 22.3G (sketch + graph) | 192s | 4 min 23 s | 7 min 35 s |
| GSearch(Densi fied MinHash) | 7.3G (sketch + graph) | 62 s | 2 min 17s | 4 min 19s |
| GSearch(Super MinHash) | 17.5G (sketch + graph) | 178s | 7 min 13 s | 10 min 54 s |
| GSearch(SetSk etch) | 14.8G (sketch + graph) | 161s | 7 min 14 s | 9 min 55 s |
| Dashing 1 | 15.4G (sketch) | ~121s | 251 min | 253 min |
| Dashing2 (setsketch) | 19.4G (sketch) | ~154s | 183 min | 186 min |
| Skani[1] | 17.6G (sketch) | - | 324 min | 324 min |
| BinDash | 16.4G (sketch) | 94s | 182 Min | 183.5 min |

[1]skani skips sparse chaining step to calculate ANI via FracMinHash for ANI < 80% comparisons, a trick can be used for other methods

**Table S5**. GSearch search (request) performance on major CPU platforms using the GTDB v207 database for graph building and 1000 genome queries. Default ProbMinHash option

| CPU | Number of threads | Clock speed (GHz) | **Request time for nt (min)** | Gene Prediction-FGSrs (min)[c] | **Request time for proteome (min)** | hmmsearch time (min)[d] | **Request time for USCG (min)** |
|---|---|---|---|---|---|---|---|
| Intel (R) Xeon (R) Gold 6226[a] | 24 | 2.70 | **2.329** | 1.348 | **1.334** | 0.524 | **0.117** |
| Intel (R) Core i7-7770HQ[b] | 8 | 2.80 | **8.654** | 6.764 | **2.041** | 1.534 | **0.510** |
| AMD EPYC 7513a[a] | 32(24 used) | 2.60 | **1.937** | 1.120 | **1.021** | 0.345 | **0.102** |
| Apple M1 Pro[b] | 10 | 3.22 | **2.369** | 2.12 | **0.866** | 0.498 | **0.168** |

[a] RHEL v7.9, Linux v3.10.0-1160, all threads used.
[b]MacOS v12.3, Darwin 21.4.0, all threads used.
[c]Parallel package was used to run multiprocess at the same time. FGSrs stands for FragGeneScanRs. Note that in practice only those genomes failing at the Request step for nucleotide-level search (best match found is <78% ANI) will be used at this step.

[d]Only 100 genomes were used for testing hmmsearch because this step is for very novel (deep-branching) genomes at order level or above, which are not very common in real-world dataset. The Parallel Package was used to run multiple processes of hmmsearch, one thread per process for hmmsearch.

**Table S6**. GSearch performance and accuracy for the viral database using the blastp-AAI as the reference standard. MASH dist command was run using 24 threads for this analysis.

| | Request (aa) | Recall (top 5) | MASH (aa) |
|---|---|---|---|
| 1000 phage | 4 min 24 s | 98.32% | 2.12 h |
| 10,000 Phage | 39 min 58 s | 96.04% | 17.4 h |

**Table S7**. Benchmarking of hnswlib-rs library using the MNIST fashion (70,000) dataset.

| | M[a] | ef_construct | recall | Build time (s) | Query time[b] (s) | Speed[c] (queries/s) |
|---|---|---|---|---|---|---|
| Run #1 | 32 | 200 | 0.9849 | 4.88 | 0.27 | 36569.63 |
| Run #2 | 64 | 400 | 0.9959 | 9.09 | 0.46 | 21699.18 |
| Run #3 | 128 | 400 | 0.9981 | 9.66 | 0.78 | 12740.01 |
| Run #4 | 128 | 800 | 0.9984 | 18.00 | 0.86 | 11574.46 |
| Run #5 | 128 | 1600 | 0.9986 | 34.53 | 0.93 | 10747.78 |
| Run #6 | 256 | 1200 | 0.9995 | 30.00 | 1.61 | 6209.15 |
| Run #7 | 256 | 1600 | 0.9996 | 39.18 | 1.67 | 5972.75 |
| Run #8 | 256 | 2400 | 0.99999 | 56.33 | 1.72 | 5804.61 |

[a] ef_search=M, the width of search for querying equals to M

[b] Large M and ef_construct take more time for both build and query search. In the MNIST fashion dataset, Euclidean distance is pre-computed and not considered in this test. The distance used is the same for Table S8, Table S9 and table S10.

[c] Tests were run on a 24 threads Intel (R) Xeon (R) Gold 6226 CPU@2.70 Ghz. Speed is tested using 10,000 queries.

**Table S8**. Benchmarking of hnswlib-rs library using the SIFT1M (1,000,000) dataset.

| | M | ef_construct | recall | Build time (min) | Query time (s) | Speed [a](queries/s) |
|---|---|---|---|---|---|---|
| Run #1 | 128 | 1600 | 0.9980 | 21.18 | 1.81 | 5528.93 |
| Run #2 | 256 | 3200 | 0.9986 | 40.83 | 3.82 | 2618.20 |

[a] Tests were run on a 24 threads Intel (R) Xeon (R) Gold 6226 CPU@2.70 Ghz. Speed is tested using 10,000 queries.

**Table S9**. Benchmarking of hnswlib-rs library for scalability using the SIFT1M dataset.

| | Number of threads | M | ef_construct | Build time (min) | Query time (s) | Speed[a] (queries/s) |
|---|---|---|---|---|---|---|
| Run #1 | 8 | 256 | 3200 | 87.97 | 7.90 | 1265.27 |
| Run #2 | 12 | 256 | 3200 | 67.02 | 6.26 | 1596.35 |
| Run #3 | 24 | 256 | 3200 | 40.83 | 3.82 | 2618.20 |

[a] Tests were run on an Intel (R) Xeon (R) Gold 6226 CPU@2.70 Ghz. For each run, the number of threads shown in column #2 were requested. Speed is tested using 10,000 queries.

**Table S10**. Comparison of hnswlib-rs with C++ hnswlib using the MNIST fashion (70,000) dataset. For this test, 60,000 genomes were used for building as database and 10,000 as query, requesting 50 best neighbors.

| | Number of threads | M | ef_construc t | Build time (s) | Query time (s) | Speed[a] (queries/s) |
|---|---|---|---|---|---|---|
| Run #1 (hnswlib-rs) | 24 | 128 | 1600 | 34.53 | 0.93 | 11225 |
| Run #2 (hnswlib-c++[1]) | 24 | 128 | 1600 | 31.02 | 1.05 | 10204 |
| Run #3 (hnswlib-rs) | 24 | 256 | 3200 | 92.83 | 2.32 | 4250 |
| Run #4 (hnswlib-c++) | 24 | 256 | 3200 | 51.83 | 1.85 | 5235 |

[1] https://github.com/nmslib/hnswlib

**Table S11**. The table shows benchmark results of FGSrs against truth (full version v0.0.1) and comparison with other tools. True positives were denoted as the number of base pair in a prediction on the correct strand, false positives the number of base pair in a prediction outside gene annotations or on the wrong strand, true negatives the number of bp outside gene annotations that weren't in any prediction, and false negatives the number of bp inside gene annotations that weren't in any prediction. Prec denotes precision, Sens denotes sensitivity, Spec denotes specificity. Negative Predictive Value and Matthew's Correlation Coefficient are provided as percentages.

| tool | TP | FP | TN | FN | prec | sens | F1 score | spec | NPV | MCC |
|---|---|---|---|---|---|---|---|---|---|---|
| FGS | 2494195 | 231257 | 656879 | 283010 | 91.51 | 89.81 | 0.90 | 73.96 | 69.89 | 62.58 |
| prodigal | 2488137 | 117535 | 745673 | 313996 | **95.49** | **88.79** | **0.92** | **86.38** | 70.37 | 70.36 |
| FGSrs | 2494195 | 231257 | 656879 | 283010 | **91.51** | **89.81** | **0.90** | **73.96** | 69.89 | 62.58 |

**Table S12**. Effect of genome completeness on GSearch recall. Genomes of different degrees of completeness were obtained by randomly sampling the gene sequences from the predicted gene collection of the complete genome. *Hydrogenimonas urashimensis*, an $H_2$ dependent chemolithoautotrophic bacteria isolated from deep sea hydrothermal vent (Mino et al., 2021), was used as query. There is no identical genome to *H. urashimensis* in the GTDB database.

| Completeness | Top Hits found/10 | Top 5 hits found/5 | Recall (top 10) | Recall (top 10) |
|---|---|---|---|---|
| 0.95 | 10/10 | 5/5 | 100% | 100% |
| 0.90 | 10/10 | 5/5 | 100% | 100% |
| 0.85 | 9/10 | 5/5 | 90% | 100% |
| 0.80 | 8/10 | 5/5 | 80% | 100% |
| 0.75 | 8/10 | 5/5 | 80% | 100% |
| 0.70 | 8/10 | 5/5 | 80% | 100% |
| 0.65 | 8/10 | 5/5 | 80% | 100% |
| 0.60 | 8/10 | 5/5 | 80% | 100% |
| 0.55 | 8/10 | 5/5 | 80% | 100% |
| 0.50 | 7/10 | 5/5 | 70% | 100% |
| 0.45 | 5/10 | 5/5 | 50% | 100% |
| 0.40 | 4/10 | 4/5 | 40% | 80% |
| 0.35 | 4/10 | 4/5 | 40% | 80% |

| | | | | |
|---|---|---|---|---|
| 0.30 | 4/10 | 4/5 | 40% | 80% |
| 0.20 | 2/10 | 2/5 | 20% | 40% |
| 0.10 | 1/10 | 2/5 | 10% | 40% |

**Table S13**. GSearch, Sourmash and Mash benchmark against blastn-ANI at nucleotide level. Query genomes is “OceanDNA-b42278.fa” from (Nishimura and Yoshizawa, 2022). Database genomes were all NCBI/RefSeq genomes. Mash, FastANI and Blastn-based ANI give the same top 10 for this query genomes. For each column, matches are ranked by the respective distance, taken directly from the software output. Top 10 found by blastn-ANI were about 80% to 97% ANI.

| Mash & FastANI & Blastn-ANI (top10), 11min for Mash, ground truth | GSearch (top10, ProbMinHash), recall 100%, 2s | Sourmash (top10), recall 90%, 13min | Dashing (top10), recall 90%, 26s | BinDash (top 10), recall 100%, 3.7s |
|---|---|---|---|---|
| GCA_902591925.1_genomic.fna | GCA_902591925.1_genomic.fna | GCA_902591925.1_genomic.fna | GCA_902591925.1_genomic.fna | GCA_902591925.1_genomic.fna |
| GCA_902617045.1_genomic.fna | GCA_902617045.1_genomic.fna | GCA_902617045.1_genomic.fna | GCA_902617045.1_genomic.fna | GCA_902617045.1_genomic.fna |
| GCA_902541175.1_genomic.fna | GCA_902547295.1_genomic.fna | GCA_902541175.1_genomic.fna | GCA_902612915.1_genomic.fna | GCA_902541175.1_genomic.fna |
| GCA_902612915.1_genomic.fna | GCA_902612915.1_genomic.fna | GCA_902612915.1_genomic.fna | GCA_902541175.1_genomic.fna | GCA_902612915.1_genomic.fna |
| GCA_004212975.1_genomic.fna | GCA_004212975.1_genomic.fna | GCA_902547295.1_genomic.fna | GCA_902547295.1_genomic.fna | GCA_902547295.1_genomic.fna |
| GCA_902547295.1_genomic.fna | GCA_902541175.1_genomic.fna | GCA_004212975.1_genomic.fna | GCA_004212975.1_genomic.fna | GCA_004212975.1_genomic.fna |
| GCA_902630885.1_genomic.fna | GCA_902586925.1_genomic.fna | GCA_902586925.1_genomic.fna | GCA_000252525.1_genomic.fna | GCA_902630885.1_genomic.fna |
| GCA_902586925.1_genomic.fna | GCA_902630885.1_genomic.fna | GCA_902631785.1_genomic.fna | GCA_902630885.1_genomic.fna | GCA_902586925.1_genomic.fna |
| GCA_902631785.1_genomic.fna | GCA_902631785.1_genomic.fna | GCA_000252525.1_genomic.fna | GCA_902631785.1_genomic.fna | GCA_902631785.1_genomic.fna |
| GCA_000252525.1_genomic.fna | GCA_000252525.1_genomic.fna | GCA_902582355.1_genomic.fna | GCA_902582355.1_genomic.fna | GCA_000252525.1_genomic.fna |

**Table S14**. GSearch (ProbMinHash), Sourmash and Dashing benchmarking against Blastn-ANI at nucleotide level. Query genome is “OceanDNA-b42278.fa” from (Nishimura and Yoshizawa, 2022). Database genomes are all NCBI/RefSeq genomes (318k) after removing the top 10 ground truth genomes found in **Table S13** above. For each column, matches are ranked by the respective distance provided in the output of each tool. Top 10 matches found by blastn-ANI showed between 75% to 80% to the query genome, corresponding to Jaccard index 0.009 to 0.015 respectively. Boldface denotes the genomes found by each method compared to the ground truth in column 1.

| Blastn-ANI (top10), ground truth | GSearch (Prob) (top10), recall 50%, 2s | Sourmash (top10), recall 60%, 14min | Dashing (default Ertl-MLE estimator) (top10), recall 10%, 26s | BinDash recall (90%). 3.7s |
|---|---|---|---|---|
| GCA_902582355.1_genomic.fna (79.24%) | **GCA_002690725.1_genomic.fna** | **GCA_002690725.1_genomic.fna** | **GCA_902582355.1_genomic.fna** | **GCA_902582355.1_genomic.fna** |
| GCA_002690725.1_genomic.fna (78.79%) | **GCA_902560315.1_genomic.fna** | **GCA_902560315.1_genomic.fna** | GCA_003282945.1_genomic.fna | **GCA_902560315.1_genomic.fna** |
| GCA_902560315.1_genomic.fna (77.92%) | **GCA_902556105.1_genomic.fna** | **GCA_902556105.1_genomic.fna** | GCA_012735275.1_genomic.fna | **GCA_002690725.1_genomic.fna** |
| GCA_902517505.1_genomic.fna (77.57%) | GCA_902558095.1_genomic.fna | **GCA_902582355.1_genomic.fna** | GCF_000165465.1_genomic.fna | **GCA_902517505.1_genomic.fna** |
| GCA_002169625.2_genomic.fna (76.41%) | **GCA_902517505.1_genomic.fna** | GCA_902558095.1_genomic.fna | GCF_003111605.1_genomic.fna | **GCA_902583575.1_genomic.fna** |
| GCA_902556105.1_genomic.fna (76.37%) | GCF_017873235.1_genomic.fna | **GCA_003213495.1_genomic.fna** | GCA_017532025.1_genomic.fna | **GCA_902579825.1_genomic.fna** |
| GCA_902583575.1_genomic.fna (75.30%) | **GCA_002169625.2_genomic.fna** | GCA_002721465.1_genomic.fna | GCA_902592715.1_genomic.fna | **GCA_003213495.1_genomic.fna** |
| GCA_003213495.1_genomic.fna (75.24%) | GCA_002704625.1_genomic.fna | **GCA_902517505.1_genomic.fna** | GCA_014653355.1_genomic.fna | **GCA_902556105.1_genomic.fna** |
| GCA_902579825.1_genomic.fna (75.15%) | GCA_902559345.1_genomic.fna | GCA_902557965.1_genomic.fna | GCA_902626385.1_genomic.fna | GCA_902558095.1_genomic.fna |
| GCA_902563835.1_genomic.fna (75.00%) | GCA_902557965.1_genomic.fna | GCA_902559345.1_genomic.fna | GCA_016288795.1_genomic.fna | **GCA_902563835.1_genomic.fna** |

**Table S15**. GSearch (SetSketch) and Dashing (Ertl’s Joint MLE) benchmarking against blastn-ANI. The table is similar to **Table S14** above but using the SetSketch algorithm.

| Blastn-ANI (top10), ground truth | GSearch with SetSketch HLL (top10), recall 60%, 2s | GSearch with SuperMinHash (top10), recall 60%, 2s | Dashing (JMLE methods, ~10x slower) (top 10), recall 10%, 17.4 min |
|---|---|---|---|
| GCA_902582355.1_genomic.fna (79.24%) | **GCA_002690725.1_genomic.fna** | **GCA_902560315.1_genomic.fna** | **GCA_902582355.1_genomic.fna** |
| GCA_002690725.1_genomic.fna (78.79%) | **GCA_902560315.1_genomic.fna** | **GCA_002690725.1_genomic.fna** | GCA_902586925.1_genomic.fna |
| GCA_902560315.1_genomic.fna (77.92%) | **GCA_902517505.1_genomic.fna** | **GCA_902583575.1_genomic.fna** | GCA_902626385.1_genomic.fna |
| GCA_902517505.1_genomic.fna (77.57%) | **GCA_902583575.1_genomic.fna** | **GCA_902517505.1_genomic.fna** | GCA_003282945.1_genomic.fna |

| | | | |
|---|---|---|---|
| GCA_002169625.2_genomic.fna (76.41%) | **GCA_902579825.1_genomic.fna** | GCA_902559345.1_genomic.fna | GCA_012735275.1_genomic.fna |
| GCA_902556105.1_genomic.fna (76.37%) | GCA_902559345.1_genomic.fna | **GCA_902579825.1_genomic.fna** | GCF_000165465.1_genomic.fna |
| GCA_902583575.1_genomic.fna (75.30%) | GCA_902558095.1_genomic.fna | **GCA_902556105.1_genomic.fna** | GCF_003111605.1_genomic.fna |
| GCA_003213495.1_genomic.fna (75.24%) | **GCA_902556105.1_genomic.fna** | GCA_902592715.1_genomic.fna | GCA_017532025.1_genomic.fna |
| GCA_902579825.1_genomic.fna (75.15%) | GCA_902528875.1_genomic.fna | GCA_014653355.1_genomic.fna | GCA_902592715.1_genomic.fna |
| GCA_902563835.1_genomic.fna (75.00%) | GCA_902551185.1_genomic.fna | GCF_000165465.1_genomic.fna | GCA_014653355.1_genomic.fna |

**Table S16**. Recall (top 10) of ProbMinHash, HyperLogLog and SuperMinHash against the blastn-ANI results.

| Mash & FastANI & Blastn-ANI (top10), 11min for Mash, ground truth | GSearch ProbMinHash (top10), recall 100%, 2s | GSearch Densified Minhash recall 100%, 5s | GSearch setsketch recall 100%, 5s | GSearch SuperMinHash recall 100%, 3s |
|---|---|---|---|---|
| GCA_902591925.1_genomic.fna | GCA_902591925.1_genomic.fna | GCA_902591925.1_genomic.fna | GCA_902591925.1_genomic.fna | GCA_902591925.1_genomic.fna |
| GCA_902617045.1_genomic.fna | GCA_902617045.1_genomic.fna | GCA_902617045.1_genomic.fna | GCA_902617045.1_genomic.fna | GCA_902617045.1_genomic.fna |
| GCA_902541175.1_genomic.fna | GCA_902547295.1_genomic.fna | GCA_902541175.1_genomic.fna | GCA_902612915.1_genomic.fna | GCA_902541175.1_genomic.fna |
| GCA_902612915.1_genomic.fna | GCA_902612915.1_genomic.fna | GCA_902612915.1_genomic.fna | GCA_902541175.1_genomic.fna | GCA_902612915.1_genomic.fna |
| GCA_004212975.1_genomic.fna | GCA_004212975.1_genomic.fna | GCA_902547295.1_genomic.fna | GCA_004212975.1_genomic.fna | GCA_004212975.1_genomic.fna |
| GCA_902547295.1_genomic.fna | GCA_902541175.1_genomic.fna | GCA_004212975.1_genomic.fna | GCA_902547295.1_genomic.fna | GCA_902547295.1_genomic.fna |
| GCA_902630885.1_genomic.fna | GCA_902586925.1_genomic.fna | GCA_902630885.1_genomic.fna | GCA_902630885.1_genomic.fna | GCA_902630885.1_genomic.fna |
| GCA_902586925.1_genomic.fna | GCA_902630885.1_genomic.fna | GCA_902586925.1_genomic.fna | GCA_902586925.1_genomic.fna | GCA_902586925.1_genomic.fna |
| GCA_902631785.1_genomic.fna | GCA_902631785.1_genomic.fna | GCA_902631785.1_genomic.fna | GCA_902631785.1_genomic.fna | GCA_902631785.1_genomic.fna |
| GCA_000252525.1_genomic.fna | GCA_000252525.1_genomic.fna | GCA_000252525.1_genomic.fna | GCA_902582355.1_genomic.fna | GCA_000252525.1_genomic.fna |

**Table S17**. Recall (top 10) of GSearch(ProbMinHash), Mash and Sourmash against the blastp-AAI results. The same query genome as shown in **Table S14** was used but at the proteome level after gene prediction. Dashing and BinDash were not included because these tools do not provide an amino acid level option.

| Blastp-AAI (top10), ground truth, AAI | GSearch ProbMinHash (top10), recall 90%, 2s | Mash, recall 90%, 13 min | Sourmash, recall 80%, 15 min |
|---|---|---|---|
| GCA_902591925.1_genomic.fna (83.2%) | **GCA_902591925.1_genomic.fna** | **GCA_902591925.1_genomic.fna** | **GCA_902591925.1_genomic.fna** |
| GCA_902617045.1_genomic.fna (83.16%) | **GCA_902617045.1_genomic.fna** | **GCA_902617045.1_genomic.fna** | **GCA_902617045.1_genomic.fna** |
| GCA_902541175.1_genomic.fna (82.57%) | **GCA_004212975.1_genomic.fna** | **GCA_902612915.1_genomic.fna** | **GCA_902541175.1_genomic.fna** |
| GCA_902612915.1_genomic.fna (82.15%) | **GCA_902612915.1_genomic.fna** | **GCA_902541175.1_genomic.fna** | **GCA_902612915.1_genomic.fna** |
| GCA_004212975.1_genomic.fna (81.86%) | **GCA_902547295.1_genomic.fna** | **GCA_902547295.1_genomic.fna** | GCA_902558095.1_genomic.fna |
| GCA_902630885.1_genomic.fna (80.99%) | **GCA_902541175.1_genomic.fna** | **GCA_004212975.1_genomic.fna** | **GCA_902547295.1_genomic.fna** |
| GCA_902547295.1_genomic.fna (80.73%) | **GCA_902586925.1_genomic.fna** | GCA_902559345.1_genomic.fna | **GCA_902630885.1_genomic.fna** |
| GCA_902586925.1_genomic.fna (80.50%) | GCA_902558095.1_genomic.fna | **GCA_902586925.1_genomic.fna** | **GCA_902586925.1_genomic.fna** |
| GCA_000252525.1_genomic.fna (79.96%) | **GCA_902630885.1_genomic.fna** | **GCA_902631785.1_genomic.fna** | **GCA_902631785.1_genomic.fna** |
| GCA_902631785.1_genomic.fna (79.48%) | **GCA_000252525.1_genomic.fna** | **GCA_902582355.1_genomic.fna** | GCA_902559345.1_genomic.fna |

**Table S18.** Running time for different values of the key parameter b and m in SetSketch based on searching 1,000 genomes against GTDB genomes (~65K).

| b | m | nt | AA | Universal |
|---|---|---|---|---|
| b=0.001 | 4096 | 3 min 11 s | 2 min 01 s | 23 s |
| b=0.0005 | 6144 | 5 min 28 s | 3 min 48 s | 39 s |
| b=0.0001 | 8192 | 13 min 19 s | 7 min 21 s | 51 s |

**Supplementary Table References**


Broder, A.Z. (1997) On the resemblance and containment of documents. In *Proceedings Compression and Complexity of SEQUENCES 1997 (Cat No 97TB100171)*: IEEE, pp. 21-29.

Christiani, T. (2020) DartMinHash: Fast Sketching for Weighted Sets. *arXiv preprint arXiv:200511547*.

Ertl, O. (2017) Superminhash-A new minwise hashing algorithm for jaccard similarity estimation. *arXiv preprint arXiv:170605698*.

Ertl, O. (2018) BagMinHash - Minwise Hashing Algorithm for Weighted Sets. *Proceedings of the 24th ACM SIGKDD International Conference on Knowledge Discovery & Data Mining*: 1368–1377.

Ertl, O. (2020) ProbMinHash – A Class of Locality-Sensitive Hash Algorithms for the (Probability) Jaccard Similarity. *IEEE Transactions on Knowledge and Data Engineering*: 1-1.

Ertl, O. (2021) SetSketch: filling the gap between MinHash and HyperLogLog. *Proc VLDB Endow* **14**: 2244–2257.

Flajolet, P., Fusy, É., Gandouet, O., and Meunier, F. (2007) Hyperloglog: the analysis of a near-optimal cardinality estimation algorithm. *Discrete Mathematics and Theoretical Computer Science*: 137-156.

Karppa, M., and Pagh, R. (2022) HyperLogLogLog: Cardinality Estimation With One Log More. *arXiv preprint arXiv:220511327*.

Mino, S., Shiotani, T., Nakagawa, S., Takai, K., and Sawabe, T. (2021) Hydrogenimonas urashimensis sp. nov., a hydrogen-oxidizing chemolithoautotroph isolated from a deep-sea hydrothermal vent in the Southern Mariana Trough. *Systematic and Applied Microbiology* **44**: 126170.

Nishimura, Y., and Yoshizawa, S. (2022) The OceanDNA MAG catalog contains over 50,000 prokaryotic genomes originated from various marine environments. *Scientific Data* **9**: 305.

Pettie, S., and Wang, D. (2022) Simpler and Better Cardinality Estimators for HyperLogLog and PCSA. *arXiv preprint arXiv:220810578*.

Shrivastava, A. (2017) Optimal densification for fast and accurate minwise hashing. *International Conference on Machine Learning*: 3154-3163.

### Supplementary Methods & Materials

#### *ProbMinHash vs. traditional MinHash*

Mash is a hashing-based algorithm based on MinHash [1], which is very efficient for comparing genome/metagenome overall similarity [2]. Mash distances represent a k-mer-based overall overlap between sequences according to a minimal evolutionary model. Essentially, Mash distance is the Jaccard similarity value of kmer shared between sequence sets A and B. However, Mash, and similar MinHash-based tools, have several limitations; most notably, the loss of k-mer frequency information (only presence/absence of kmer is counted) and the impact of relative set size (e.g., completeness level of a genome) on the Jaccard similarity estimates (for example, Mash distance, where MinHash based estimation of Jaccard index is biased by different set size, or total k-mer count of the genome due to bottom-k sketches, HyperLogLog is not affected by different set size) [2, 3] Although some recent MinHash implementations address the relative set size limitation (e.g., the over-sketching and track-abundance methods of the MinHash-based tools 'finch , 'sourmash' or FracMinHash (sketching a subset of k-mers according to size of set) and HyperLogLog) [4-7], they do not utilize the frequencies of all observed k-mers in generating the k-mer-profile (sketch) for a given sequence set. More recently, in the HULK software, consistent weighted sampling ($D^2$histosketch which is the same with P-MinHash algorithm and it was proposed for $J_p$, we called it P-MinHash because it was invented before $D^2$histosketch and is equivalent to P-MinHash[8]) [9] was utilized to incorporate k-mer frequency information when estimating weighted and standard Jaccard similarity, which effectively addresses these limitations mentioned above [10]. Notably, the hash algorithm

(P-MinHash) used in $D^2$histosketch could be further optimized to achieve a time complexity below *O(nm)* (where m denotes the signature size and n is the number of elements with nonzero weight in two sequence sets), further improving the performance of applications such as HULK. Motivated by the SuperMinHash for conventional Jaccard similarity estimation [11] and BagMinHash algorithm for weighted Jaccard similarity estimation [12], ProbMinHash (probminhash 3(a) and 4 algorithm) is orders of magnitude faster than the original algorithm P-MinHash proposed in $D^2$hist[13]osketch [14]. Probminhash estimates the Jaccard probability $J_p$ index, and 1- Jp is indeed a metric on probability distributions and is Pareto optimal (Supplementary Note 1) [8, 14]. Densified MinHash, or One Permutation MinHash with Optimal/faster Densification, is the fastest MinHash algorithm due to theoretical breakthrough (average case O(n+m)) despite large variance than classic MinHash (Table S2). It uses only one hash function but copy values from empty bins to non-empty bins ("densified") in the sketch vector either mapping forward or backward or both [13, 15, 16]. We choose Sourmash, Mash, Dashing 1/2 and BinDash for benchmark because all are estimation of Jaccard index, which correlates very well with ANI after transformation and were all benchmarked against blastn-based ANI and fastANI [2, 19].

**Comparison with other genome/sequence search algorithm**

There are many other data structures (designed for general purposes sequence search problems) such as Sequence Bloom Tree (SBT) and its variants [20-22] , COBS/BIGSI [23, 24], Layered LSH (approximating ungapped alignment, not applicable to ANI like distance, which is gapped alignment) [25] and RAMBO (Repeated and Merged Bloom Filter) [26] that can achieve linear or sometimes even sub-linear genomic database search performance alone. However, those tools have never been benchmarked against blast-based ANI/AAI in the context of microbial genomic search (e.g., that is whether the best hits/genomes found by the mentioned tools are the same with ANI/AAI comparison best hits/genomes), which is the golden standard for measuring microbial genomic distance/identity and thus infer microbial taxonomy [27, 28]. MinHash-based tool, however, such as Mash and FastANI are based on evolutionary model and were benchmarked against Blastn-based ANI and were thought to be the most accurate k-mer-based sub-linear algorithms for ANI comparisons and/or searching [2, 19].

***HNSW in Rust benchmark against testing dataset***

To benchmark our reimplementation of hnswlib, we followed standard ANN benchmark procedures using two popular testing datasets (MINST and SIFT1M) based on their Euclidean distance[29]. Our results showed that, for the MINST fashion dataset (784 dimensions, 60,000 vectors), recall for top 100 neighbors of 10,000 query vectors is greater than 98% for a smaller number of M and ef_construct, and even higher recall rate (99.86%) for a medium M and ef_construct while query speed is not compromised (Supplemental Table S7). For the SIFT1M dataset (128 dimensions, 1,000,000 vectors), recall for top 100 neighbors of 10,000 query vectors was 99.77% for a medium M and ef_construct (Supplemental Table S7 and S8). In terms of speed, we compare it with hnswlib using the MINST-fashion dataset and it is as fast as hnswlib: it took 18.06s and

0.89s for database building and searching for hnswlib-rs while it took 18.47s and 1.07s for database building and searching for the C++ hnswlib) (Supplementary Table S9). The Rust package hnswlib-rs can be found at: https://github.com/jean-pierreBoth/hnswlib-rs. For each genomic database, we chose M and ef_construct experimentally, by gradually increasing M and ef_construct while monitoring query speed and recall, similar to what is shown in Supplementary Table S2 for MNIST dataset. We stopped the assessment when there was only a marginal increase in accuracy but decent decrease in speed. To leverage between recall and speed, we use M=128 and ef_search=1600 for graph building for GTDB database fungal database while M=128, ef_search=3200 for phage database.

***Details of SetSketch implementation***

We implemented SetSketch algorithm 1 locality sensitivity section (LSH) according to lower and upper bound of Jaccard ($J_{low} = max(0, \frac{b^{\frac{D_0/m+1}{2}} - 1}{b-1} - 1), J_{up} = \frac{b^{D_0/m} - 1}{b-1}$) where $D_0$ is the number of registers in the sketch of genome A that are equal to those in the sketch of genome B. In practice, $J_{low}$ is closer to true Jaccard for small J thus we use it. We use parameter $m$ = 6144, b = 1.0005, a = 20, and q = 65534 instead of the default ones to have smaller RMSE (<0.8%) around J=0.015 (corresponding to ANI 77.99% according to Mash equation) and acceptable running time (Figure S8, Table S18), which is equivalent to HyperLogLog for estimating Jaccard index in terms of space but with smaller variance (with b close to 2, SetSketch will then be equivalent to SuperMinHash). A SetSketch using this configuration is suitable to represent any set with up to 10^19 distinct elements/k-mers (much larger than total-number of k-mers from microbial genomes). The expected error of cardinality estimates or LSH is very small ($\sqrt{\frac{1}{m}(\frac{b+1}{b-1}log(b)-1)}$) as b approximates 1($\frac{1}{\sqrt{m}}$) [30], close to that of MinHash for large m, like 10^4 or above but use much smaller space. We also implemented the Joint Maximum Likelihood Estimator (JMLE): the ML estimate for Jaccard was found by standard univariate optimization algorithm called Brent's method, based on the argmin Rust package since the ML function is strictly concave for the parameter mentioned above [30]. The RMSE of JMLE based on its Fisher information can be found in Supplementary Note 7. Since RMSE of JMLE is smaller than LSH, it is much slower in practice, but we only use it for filtering false positives after top k best neighbors were found for each query by LSH and HNSW, thus not a problem for overall speed.

**Details of Densified MinHash implementation**

We reimplemented both One Permutation MinHash with Optimal Densification [15] and also Faster Densification [16] in Rust in the probminhash package. Specifically, one predefined hash function will be applied to all elements/k-mers in the set. Then, one-permutation MinHash deterministically partitions the hash values into a predefined maximum number B of buckets, extracts the smallest hash value in each bucket, and extracts the b lowest bits of each smallest hash value. These B  b bits are used as the signature of the set. Usually, a hash value v is assigned to the ⌈v/(M/B)⌉ bucket, where M is the maximum possible value for v.  Although fast at both constructing and comparing sketches, one-permutation MinHash may produce a bucket that contains hash values for one set but no hash values for another set.  All buckets are totally ordered in some way, each empty bucket uses the smallest hash value in the next non-empty bucket as its own hash value (densified), and an additional bucket containing a special value is ordered after all other buckets. We either mapping non-empty bins to empty bins (faster densification) or the other way around to copy values (optimal densification). The densified sketch vectors for 2 sets were then used for calculating collision probability/Jaccard index (See densminhash.rs in prominhash package). We use 10 times larger sketch size m for densified MinHash to achieve a smaller variance compared to original MinHash.

#### Supplementary Methods & Materials References


1. Broder, A.Z. in Proceedings. Compression and Complexity of SEQUENCES 1997 (Cat. No. 97TB100171) 21-29 (IEEE, 1997).
2. Ondov, B.D. et al. Mash: fast genome and metagenome distance estimation using MinHash. *Genome Biology* **17**, 132 (2016).
3. Koslicki, D. & Zabeti, H. Improving MinHash via the containment index with applications to metagenomic analysis. *Applied Mathematics and Computation* **354**, 206-215 (2019).
4. Brown, C.T. & Irber, L. sourmash: a library for MinHash sketching of DNA. *Journal of Open Source Software* **1**, 27 (2016).
5. Bovee, R. & Greenfield, N. Finch: a tool adding dynamic abundance filtering to genomic MinHashing. *Journal of Open Source Software* **3**, 505 (2018).
6. Irber, L. et al. Lightweight compositional analysis of metagenomes with FracMinHash and minimum metagenome covers. *bioRxiv*, 2022.2001.2011.475838 (2022).
7. Baker, D.N. & Langmead, B. Dashing: fast and accurate genomic distances with HyperLogLog. *Genome Biology* **20**, 265 (2019).
8. Moulton, R. & Jiang, Y. Maximally Consistent Sampling and the Jaccard Index of Probability Distributions. *2018 IEEE International Conference on Data Mining (ICDM)*, 347-356 (2018).
9. Yang, D., Li, B., Rettig, L. & Cudré-Mauroux, P. $D^2$histoSketch: Discriminative and Dynamic Similarity-Preserving Sketching of Streaming Histograms. *IEEE Transactions on Knowledge and Data Engineering* **31**, 1898-1911 (2019).
10. Rowe, W.P.M. et al. Streaming histogram sketching for rapid microbiome analytics. *Microbiome* **7**, 40 (2019).
11. Ertl, O. Superminhash-A new minwise hashing algorithm for jaccard similarity estimation. *arXiv preprint arXiv:1706.05698* (2017).
12. Ertl, O. BagMinHash - Minwise Hashing Algorithm for Weighted Sets. *Proceedings of the 24th ACM SIGKDD International Conference on Knowledge Discovery & Data Mining*, 1368–1377 (2018).
13. Jia, P. et al. in Proceedings of the 2021 International Conference on Management of Data 830-842 (2021).
14. Ertl, O. ProbMinHash – A Class of Locality-Sensitive Hash Algorithms for the (Probability) Jaccard Similarity. *IEEE Transactions on Knowledge and Data Engineering*, 1-1 (2020).

15. Shrivastava, A. Optimal densification for fast and accurate minwise hashing. *International Conference on Machine Learning*, 3154-3163 (2017).
16. Mai, T. et al. in Uncertainty in Artificial Intelligence 831-840 (PMLR, 2020).
17. Baker, D.N. & Langmead, B. Dashing 2: genomic sketching with multiplicities and locality-sensitive hashing. *bioRxiv* (2022).
18. Agret, C., Cazaux, B. & Limasset, A. Toward optimal fingerprint indexing for large scale genomics. *bioRxiv*, 2021.2011.2004.467355 (2022).
19. Jain, C., Rodriguez-R, L.M., Phillippy, A.M., Konstantinidis, K.T. & Aluru, S. High throughput ANI analysis of 90K prokaryotic genomes reveals clear species boundaries. *Nature Communications* **9**, 5114 (2018).
20. Solomon, B. & Kingsford, C. Fast search of thousands of short-read sequencing experiments. *Nature biotechnology* **34**, 300-302 (2016).
21. Solomon, B. & Kingsford, C. Improved search of large transcriptomic sequencing databases using split sequence bloom trees. *Journal of Computational Biology* **25**, 755-765 (2018).
22. Sun, C., Harris, R.S., Chikhi, R. & Medvedev, P. Allsome sequence bloom trees. *Journal of Computational Biology* **25**, 467-479 (2018).
23. Bingmann, T., Bradley, P., Gauger, F. & Iqbal, Z. in String Processing and Information Retrieval: 26th International Symposium, SPIRE 2019, Segovia, Spain, October 7–9, 2019, Proceedings 26 285-303 (Springer, 2019).
24. Bradley, P., Den Bakker, H.C., Rocha, E.P., McVean, G. & Iqbal, Z. Ultrafast search of all deposited bacterial and viral genomic data. *Nature biotechnology* **37**, 152-159 (2019).
25. Chakraborty, A. & Bandyopadhyay, S. in 2018 Fifth International Conference on Emerging Applications of Information Technology (EAIT) 1-4 (IEEE, 2018).
26. Gupta, G. et al. in Proceedings of the 2021 International Conference on Management of Data 2226-2234 (2021).
27. Konstantinidis, K.T. & Tiedje, J.M. Towards a Genome-Based Taxonomy for Prokaryotes. *Journal of Bacteriology* **187**, 6258-6264 (2005).
28. Goris, J. et al. DNA–DNA hybridization values and their relationship to whole-genome sequence similarities. *International Journal of Systematic and Evolutionary Microbiology* **57**, 81-91 (2007).
29. Aumüller, M., Bernhardsson, E. & Faithfull, A. ANN-Benchmarks: A benchmarking tool for approximate nearest neighbor algorithms. *Information Systems* **87**, 101374 (2020).
30. Ertl, O. SetSketch: filling the gap between MinHash and HyperLogLog. *Proc. VLDB Endow.* **14**, 2244–2257 (2021).

### Supplementary Notes

**Note 1. Calculation of metric (Weighted) Jaccard similarity indices.** $J_W = \frac{\sum_{d\in D} min(\omega_A(d),\omega_B(d))}{\sum_{d\in D} max(\omega_A(d),\omega_B(d))}$ (where $\omega$ is the weight function of each element $d$) and total k-mer count in genomes in biasing the Jaccard index estimation (or set size) [1]. K-mer-weighted hashing approaches (taking into account the abundance of k-mers, not just presence/absence) are more space-expensive (e.g. higher memory requirement) for large dataset [2], but are advantageous for genomes with frequent repeats. They have not been widely adopted yet [3, 4]. To consider multiplicity of k-mers in the k-mer set of genomes, traditional MinHash algorithms will not be a good choice since they assume unique set element (k-mer). New MinHash algorithms such as ICWS, BagMinHash and DartMinHash were designed for weighted set to address this limitation, with DartMinHash being the fastest [5-8]. Still, those weighted MinHash algorithms do not solve the problem of different genome size (set size) in biasing estimation of weighted Jaccard index [9]. A possible solution is to normalize the abundance of k-mer by the total k-mer count of each dataset, threby providing a probability distribution of each k-mer. This then leads to the normalized weighted Jaccard index $J_N = \frac{\sum_{d\in D} min(\frac{\omega_A(d)}{\sum_{d'\in D}\omega_A(d')},\frac{\omega_B(d)}{\sum_{d'\in D}\omega_B(d')})}{\sum_{d\in D} max(\frac{\omega_A(d)}{\sum_{d'\in D}\omega_A(d')},\frac{\omega_B(d)}{\sum_{d'\in D}\omega_B(d')})}$, where $\omega_A$and $\omega_B$ describe the weight of each k-mer. Also, those weighted MinHash algorithm could be further optimized computationally to be orders of magnitude faster, like it was done in BagMinHash and DartMinHash [10], similar to the computational optimizations implemented in MinHash in SuperMinHash [11]. Recently, ProbMinHash was proposed to take into account both weighted set (k-mer multiplicity) and total set size (total k-mer count, or genome size) [12]. Accordingly, new Locality Sensitive Hashing algorithms (P-MinHash) considering weighted set and different set size was proposed to estimate weighted and normalized (to account for set size difference) Jaccard-like index $J_P = \sum_{d \in D} \frac{1}{\sum_{d' \in D} max(\frac{\omega_A(d')}{\omega_A(d)}, \frac{\omega_B(d')}{\omega_B(d)})}$. P-MinHash is considered a more general case for Jaccard Index estimation, and is more close to true Jaccard than $J_W$ and $J_N$ [13, 14]. More importantly, $J_p$ is Pareto optimal and 1-$J_p$ (similarity) is a proper metric [13]. It has been shown that ANI estimated from $J_p$, which was computed by ProbMinHash (ProbMinHash2 algorithm) in Dashing 2, is slight better for bacterial genomes than traditional MinHash like Mash [15]. ProbMinHash was built upon the new MinHash algorithm with further computational optimization for speed [12]. Hence, ProbMinHash is the currently the default optioin in GSearch. HyperLogLog can also be used to estimate Jaccard index: distinct element count of k-mers in set/genome A and B via HyperLogLog sketch (memory efficiency) and then use the inclusion-exclusion rule to have Jaccard index ($Jaccard(A, B) = \frac{|A|+|B|-|A \cup B|}{|A \cup B|}$) as implemented in Dashing, despite being overall less accurate for small cardinalities (e.g. virus genomes) [16, 17]. The Maximum Likelihood estimator (MLE) [16], in addition to the original estimator in HyperLogLog and improved estimator in [16] for distinct element counting, was thought to have the smallest variance, which met the Cramér-Rao lower bound, despite slower and difficult to compute and update [18]. This is the idea behind Dashing, which we benchmark GSearch against.

**Note 2. Kmer selecting rationale for nucleotide (nt) and amino-acid (aa) level searches for MinHash/probminhash tools**. Assume amino acid/nucleotide sequences evolve at a constant rate and alphabets $|\Sigma|$ for AA is 20 and 4 for nucleotide. Consider two sequences x , y $\in A^N$ drawn randomly over the alphabet of size #A = 20, N is the genome length, and let $v_x, v_y \in R^{20^k}$ denote the k-mer frequency profile (multiplicity of each k-mer divided by total number of k-mers) for each sequence. For long sequences with N ≫ $20^k$, k-mer frequencies will converge to their mean, that is $20^{-k}$ for all their components, which implies that $\|v_x - v_y\|_1 \rightarrow 0$. Therefore, any k-mer profile-based method will severely underestimate distance between these two random sequences. In order to avoid this, k has to be restricted to large values $k' \geq log_{|\Sigma|}(N)$. In practice, since genomes sequences are not completely random, k can be slightly smaller than $\log_4(N)$ without underestimating genomic distance. Ondov et al., in the MASH paper, came up with a probability term (1-q)/q to take into account the probability of having a random kmer: $k' \geq log_{|\Sigma|}(N(1-q)/q)$ [19]. Assume a probability q=0.01, for a typical bacterial proteome, k>=$\log_{20}$(1000000*0.99/0.01) =6.146, so at least a kmer of size 6 or above should be used. For a typical bacterial genome at nt level, N is about 4*10^6, k >= $\log_4$(4*10^6*0.99/0.01) =14.280, so at least a kmer of size 14 or above should be used. For universal genes only, N is about 350 AA * 120=42000, so k >= $\log_{20}$(42000*0.99/0.01) = 5.087, so at least a kmer of size 5 or above should be used. Since universal gene alphabets are not randomly evolving (e.g., some regions in genes that encode key metabolic functions such as RNA processing units rarely mutate), k=5 should be appropriate. To optimize between sensitivity and specificity for kmer, we followed the practice suggested by Ondov et al., and Jain et al. [19, 20]; that is, use k=16 for bacterial nucleotide genome sequences and a k=7 for bacterial proteome sequences. For fungal genomes, genome size is 10 to 20 times larger than tht of bacteria but the same formula applies and so, we have k >= $\log_4$(20*4*10^6*0.99/0.01) =16.441, and accordingly we use kmer=21. For fungal proteome, coding density is much smaller than that of bacteria (we use an empirical 0.5 coding density) thus, we have k >= $\log_{20}$(10*1*10^6*0.99/0.01) =6.914. We use k=11 for fungal proteome to increase the specificity of the searches. For virus/bacteriophage genomes, we use k=11 for nt and k=7 for aa, considering also the jumbo/giant phage genomes that were found recently.

**Note 3. Assessment of the time complexity of ProbMinHash and HNSW**. The search time complexity is estimated based on the equation (or big O annotation) $O(vdlog(n))$, where n is the dataset size while v and d are maximum out-degree of the graph to be built and the number of dimensions (kmer features in a genome) of the dataset, respectively [21, 22]. In nearest neighbor search studies, L2 distance (Euclidean distance) is often used and is related to dataset dimension, which is very large for k-mer features of genomes (~10$^{\wedge 6}$). In the case of ProbMinHash distance, which is a MinHash-based method to sample the k-mer space of genomes and serve as a dimensionality reduction in approximating genomic distance, the sampled number of k-mers (or meanwise hashes) is always a constant and much smaller number (sketch size, normally around 10,000) than the total number of k-mers of a genome (number of dimensions of dataset).

Therefore, d can be ignored (MinHash samples only a small fraction of k-mer space/dimension). Also, v is a small constant considering the graph structure to be built (normally smaller than 100). Therefore, the time complexity, as it was also explained in the main text, can be safely written as $O(log(n))$ for low dimension datasets [22]. Similar rules applied for the time complexity of the graph build step, which is $O(dnlog(n))$ and thus, $O(nlog(n))$ in our case, where d is dataset dimension. Time complexity $O(n + mlog(m))$ of the Probminhash3a algorithm, where n is the set size (total sampled k-mer number) while m is the number of weighted elements (k-mers), is also a constant in the context of high-quality genomes given also that m is very small for prokaryotic and bacteriophage genomes (normally about 5% genomic sequences are multi-copy). Therefore, ProbMinHash time complexity is very close to $O(n)$ in practice. For a given genome, for example, bacterial genome or viral genome, n is the sampled k-mer number of the genome, and thus also a constant number.

**Note 4. Kmer-based genomic distance estimation is less accurate for distantly related genomes**. For a random mutation rate $r \in (0,1)$, the probability that a k-mer (k Is length of the k-mer) belonging to sequence X and Y is not mutated is $1 - (1 - r)^k \approx 1 - exp(-\frac{r}{k})$ , indicating that $k \lesssim r^{-1}$, so k must be small enough to capture the mutation and also big enough to avoid underestimation (**Note 2**). This means that for close related bacterial genomes (r<0.01 for example), k could easily meet both conditions mentioned above while for distantly related genomes -for example- r=0.125

(corresponding to an ANI value of 87.5%) k <= 8, which is contradictory with $k > \log_4(N) = 14.28$ (Note 2). Thus, k-mer based method will lose accuracy for distantly related genomes. In practice, we observed that this r threshold is around 0.215 (ANI=78.5%) because mutation is not completely random as assumed above.

**Note 5. Theoretical guarantee of graph based NNS search algorithms**. Until recently, a theoretical analysis based on a dataset evenly distributed on an d-dimension Euclidean sphere ($d \ll \log(n)$, n is the dataset size) showed that under certain conditions, there is a guarantee that the best neighbors could be found compare to brute-force distance metric comparisons [23]. However, a theoretical analysis under more general conditions, e.g., other metric space, or d not being much smaller than log(n) is still not available, to the best of our knowledge. Despite the lack of theoretical analysis under more general conditions, graph-based algorithms work well in practice, as also reflected by the large number of graph-based NNS libraries available [21, 24].

**Note 6. Parallelism of probminhash and HNSW**. Due to ownership mechanisms of Rust (so called memory and thread safety), fearless concurrency (e.g., no data competition/race, memory and thread safety) is made possible. The crossbeam crate package was used in GSearch for communication of data among threads for task level parallelism while Rayon crate was used to do data level parallelism [25]. Accordingly, by default, GSearch uses all available processors/threads to make full use of multiprocessors of host machine.

**Note 7. Details of Joint Maximum Likelihood estimator in SetSketch algorithm**. For cardinalities $n_U$ and $n_v$ of set U and V estimated by SetSketch cardinality estimation section according to equation 12 in [26], the maximum likelihood method can be used to estimate J. Specifically, the log-likelihood function as a function of J is: $log\mathcal{L}(J) = D_+ log(p_b(u - vJ)) + D_- log(p_b(v - uJ)) + D_0 log(1 - p_b(u - vJ) - p_b(v - uJ))$, where $D_+ = |\{i: K_{Ui} > K_{Vi}\}|, D_- = |\{i: K_{Ui} < K_{Vi}\}|, D_0 = |\{i: K_{Ui} = K_{Vi}\}|$ are number of registers in the sketch of U that are greater than, less than, or equal to those in the sketch of V, respectively; $K_{Ui}$ $and$ $K_{Vi}$ are registers of set U and V, respectively while v and u is relative cardinalities $u = \frac{n_U}{n_U + n_V}$ and $v = \frac{n_v}{n_U + n_V}$, respectively. $p_b(x)$ is defined as $p_b(x) = -log_b(1 - x\frac{b-1}{b})$. The RMSE of the ML estimate is expected to be $I^{-1/2}(J)$, where I denotes the Fisher information with respect to J for $n_U$ and $n_v$:

$$I(J) = \frac{m(b-1)^2}{b^2 log^2(b)}\left(\frac{(vb^{p_b(u-vJ)})^2}{p_b(u-vJ)} + \frac{(ub^{p_b(v-uJ)})^2}{p_b(v-uJ)} + \frac{(vb^{p_b(u-vJ)} + ub^{p_b(v-uJ)})^2}{1 - p_b(u-vJ) - p_b(v-uJ)}\right)$$

The SetSketch paper showed that the estimation error (RMSE) for J will be almost the same for MinHash with the same number of registers m.

#### Supplementary Notes References

6. Shrivastava, A. Simple and efficient weighted minwise hashing. *Advances in Neural Information Processing Systems* **29** (2016).
7. Christiani, T. DartMinHash: Fast Sketching for Weighted Sets. *arXiv preprint arXiv:2005.11547* (2020).
8. Wu, W., Li, B., Chen, L., Gao, J. & Zhang, C. A review for weighted minhash algorithms. *IEEE Transactions on Knowledge and Data Engineering* **34**, 2553-2573 (2020).
9. Koslicki, D. & Zabeti, H. Improving MinHash via the containment index with applications to metagenomic analysis. *Applied Mathematics and Computation* **354**, 206-215 (2019).
10. Ertl, O. in Proceedings of the 24th ACM SIGKDD International Conference on Knowledge Discovery & Data Mining 1368–1377 (Association for Computing Machinery, London, United Kingdom; 2018).
11. Ertl, O. Superminhash-A new minwise hashing algorithm for jaccard similarity estimation. *arXiv preprint arXiv:1706.05698* (2017).
12. Ertl, O. ProbMinHash – A Class of Locality-Sensitive Hash Algorithms for the (Probability) Jaccard Similarity. *IEEE Transactions on Knowledge and Data Engineering*, 1-1 (2020).
13. Moulton, R. & Jiang, Y. in 2018 IEEE International Conference on Data Mining (ICDM) 347-356 (2018).
14. Yang, D., Li, B., Rettig, L. & Cudré-Mauroux, P. $D^2$histoSketch: Discriminative and Dynamic Similarity-Preserving Sketching of Streaming Histograms. *IEEE Transactions on Knowledge and Data Engineering* **31**, 1898-1911 (2019).
15. Baker, D.N. & Langmead, B. Dashing 2: genomic sketching with multiplicities and locality-sensitive hashing. *bioRxiv* (2022).
16. Ertl, O. New cardinality estimation algorithms for HyperLogLog sketches. *arXiv preprint arXiv:1702.01284* (2017).
17. Baker, D.N. & Langmead, B. Dashing: fast and accurate genomic distances with HyperLogLog. *Genome Biology* **20**, 265 (2019).
18. Pettie, S. & Wang, D. Simpler and Better Cardinality Estimators for HyperLogLog and PCSA. *arXiv preprint arXiv:2208.10578* (2022).
19. Ondov, B.D. et al. Mash: fast genome and metagenome distance estimation using MinHash. *Genome Biology* **17**, 132 (2016).
20. Jain, C., Rodriguez-R, L.M., Phillippy, A.M., Konstantinidis, K.T. & Aluru, S. High throughput ANI analysis of 90K prokaryotic genomes reveals clear species boundaries. *Nature Communications* **9**, 5114 (2018).
21. Lu, K., Kudo, M., Xiao, C. & Ishikawa, Y. HVS: hierarchical graph structure based on voronoi diagrams for solving approximate nearest neighbor search. *Proceedings of the VLDB Endowment* **15**, 246-258 (2021).
22. Malkov, Y.A. & Yashunin, D.A. Efficient and Robust Approximate Nearest Neighbor Search Using Hierarchical Navigable Small World Graphs. *IEEE Transactions on Pattern Analysis and Machine Intelligence* **42**, 824-836 (2020).
23. Prokhorenkova, L. & Shekhovtsov, A. Graph-based Nearest Neighbor Search: From Practice to Theory. *Proceedings of the 37th International Conference on Machine Learning* **119**, 7803--7813 (2020).
24. Fu, C., Xiang, C., Wang, C. & Cai, D. Fast approximate nearest neighbor search with the navigating spreading-out graph. *arXiv preprint arXiv:1707.00143* (2017).
25. Moraza, I.E. Rust High Performance. (Packt Publishing, Birmingham, UK; 2018).
26. Ertl, O. SetSketch: filling the gap between MinHash and HyperLogLog. *Proc. VLDB Endow.* **14**, 2244–2257 (2021).
